# Modeling cellular crosstalk and organotypic vasculature development with human iPSC-derived endothelial cells and cardiomyocytes

**DOI:** 10.1101/2020.05.04.075846

**Authors:** Emmi Helle, Minna Ampuja, Alexandra Dainis, Laura Antola, Elina Temmes, Eero Mervaala, Riikka Kivelä

## Abstract

**Rationale:** Cell-cell interactions are crucial for the development and function of the organs. Endothelial cells act as essential regulators of tissue growth and regeneration. In the heart, endothelial cells engage in delicate bidirectional communication with cardiomyocytes. The mechanisms and mediators of this crosstalk are still poorly known. Furthermore, endothelial cells in vivo are exposed to blood flow and their phenotype is greatly affected by shear stress.

**Objective:** We aimed to elucidate how cardiomyocytes regulate the development of organotypic phenotype in endothelial cells. In addition, the effects of flow-induced shear stress on endothelial cell phenotype were studied.

**Methods and results:** Human induced pluripotent stem cell (hiPSC) -derived cardiomyocytes and endothelial cells were grown either as a monoculture or as a coculture. hiPS-endothelial cells were exposed to flow using the Ibidi-pump system. Single-cell RNA sequencing was performed to define cell populations and to uncover the effects on their transcriptomic phenotypes. The hiPS-cardiomyocyte differentiation resulted in two distinct populations; atrial and ventricular. Coculture had a more pronounced effect on hiPS-endothelial cells compared to hiPS-cardiomyocytes. Coculture increased hiPS-endothelial cell expression of transcripts related to vascular development and maturation, cardiac development, and the expression of cardiac endothelial cell -specific genes. Exposure to flow significantly reprogrammed the hiPS-endothelial cell transcriptome, and surprisingly, promoted the appearance of both venous and arterial clusters.

**Conclusions:** Single-cell RNA sequencing revealed distinct atrial and ventricular cell populations in hiPS-cardiomyocytes, and arterial and venous-like cell populations in flow exposed hiPS-endothelial cells. hiPS-endothelial cells acquired cardiac endothelial cell identity in coculture. Our study demonstrated that hiPS-cardiomoycytes and hiPS-endothelial cells readily adapt to coculture and flow in a consistent and relevant manner, indicating that the methods used represent improved physiological cell culturing conditions that potentially are more relevant in disease modelling. In addition, novel cardiomyocyte-endothelial cell crosstalk mediators were revealed.

## Introduction

Endothelial cells (ECs) line the interior surfaces of blood vessels throughout the whole body. ECs are metabolically active, control vasomotor tone, and regulate angiogenesis. *In vivo,* endothelial cells acquire specific identities according to diverse stimuli from blood flow, hormones, and crosstalk signals from the nearby parenchymal cells ^1–3^. Arterial, venous, and lymphatic ECs exhibit their specific gene expression profiles, as do ECs in different organs in the body. The organotypic heterogeneity of endothelial cells is now considered as their core property, but the mechanisms regulating the organotypic features are still largely unknown. Isolated ECs, however, have been demonstrated to lose their tissue identity within a couple of days in *in vitro* culture after isolation, highlighting the importance of the niche signaling ^4,5^.

ECs comprise a significant proportion of the cells in the heart ^6^. Endothelial cell-cardiomyocyte crosstalk is an important process during cardiac development, but also in adult life ^7^. Cardiomyocytes (CMs) produce and secrete numerous proteins and peptides, called cardiokines or cardiomyokines. For signaling with ECs, angiogenic factors such as VEGFs, FGFs, HGF, and angiopoietin 1 are among the most important ones ^8^. The cardiac EC secretome, in turn, includes angiocrines such as endothelin-1 and nitric oxide that are known to regulate contractile responses of CMs ^9,10^, and apelin and neuregulin-1, which regulate CM proliferation and growth ^11,12^.

Since the method of deriving induced pluripotent stem cells (hhiPSCs) from adult somatic cells was developed ^13^, numerous different cell types have been differentiated from these cells, including ECs and CMs. hiPSCs are a powerful tool to study specific diseases in patient-derived cells. Many established models using hiPSCs have shown that patient disease phenotypes can be recapitulated in vitro ^14^. However, hiPS-EC differentiation protocols often result in a cell population expressing immature EC markers, potentially resulting in plasticity to further develop to, or even transfer between arterial and venous phenotypes ^15,16^. Similarly, most hiPS-CM differentiation protocols result in an immature cell population with an embryonic-like gene expression pattern ^17^. Although with time the cells mature, at a given time point hiPS-CMs likely represent a heterogenous mixture of different cardiac subtypes and maturation stages.

Our aim was to examine the crosstalk between hiPS-ECs and hiPS-CMs by studying how the transcriptomic profiles of these cells are changed by the presence of the other cell type. In addition, the role of shear stress in modifying the hiPS-ECs phenotype was studied. Our hypothesis was that both coculture and flow result in hiPS-ECs and hiPS-CMs that are more relevant for cardiac disease modeling. We demonstrate that ECs acquire properties towards cardiac-specific ECs and increase the expression of genes related to cardiovascular development in coculture. Both coculture and flow improved the maturity and quiescence of the ECs. Our results indicate that flow and coculture provide more physiological cell culture conditions for hiPS-ECs, resembling those *in vivo*.

## Methods

An expanded Methods section is available in the Online Data Supplement.

### hiPS cell lines

Three human induced pluripotent stem cell lines (HEL47.2, HEL46.11, HEL24.3) were purchased from Bio-medicum Stem Cell Cen-ter Core Facility. The cell lines were created by using retroviral/Sendai virus transduction of Oct3/4, Sox2, Klf4, and c-Myc, as described previously^18–20^. In addition, hiPS line K1 was a kind gift from Prof. Anu Wartiovaara group.

### hiPSC culture and differentiation

Detailed information of hiPSC culture, hiPS-EC differentiation and hiPS-CM differentiation is presented in the Online Data Supplement. hiPS-CM HEL47.2 cells were 41 days old, and HEL24.3 cells were 37 days old at the time of cell collection for single-cell RNA-seq.

The following methods used for cell characterization are presented in detail in the Online Data Supplement: qPCR analysis on CM marker gene expression during hiPS-CM differentiation, immunofluorescence staining of cells, Matrigel tube formation assay, and LDL uptake assay.

### Optogenetic analysis of hiPC-CMs electrophysiology

For optogenetic analysis, lentiviral vector Optopatch (a kind gift from Adam E. Cohen group ^21,22^ acquired through Addgene, plasmid # 62984), was introduced in hiPS-CMs. Action potentials were recorded from spontaneously beating hiPS-CMs expressing Optopatch. Data was normalized by fitting the acquired signal to an exponential function with cPot Cardiac Action Potential Calculator software, written in MATLAB. More detailed information on the experiment is presented in the Online Data Supplement. In addition, the optogenetic imaging platform and workflow is described in more detail in Björk et al., 2017 ^23^.

### hiPS-EC exposure to flow

After sorting, 2.5-3.5 x 10^5 hiPS-ECs were plated on an Ibidi µ-Slide I Luer (80176, Ibidi). Next day, the cells were subjected to laminar shear stress of 15 dyn by using the Ibidi Pump System (10902, Ibidi). After 24 hours of exposure to flow, the cells were processed either for RNA-sequencing or single-cell RNA-sequencing. For RNA-sequencing, the cells were collected into the RA1 lysis buffer and extracted using the Nucleospin RNA Plus Extraction kit (740984, Macherey-Nagel).

### hiPS-EC and hiPS-CM coculture

For coculture, hiPS-CMs and hiPS-ECs were plated together in BPEL media with 50 ng/ml VEGF. The concentration of cells was calculated using Bio-Rad TC10 or TC20 Automated Cell Counter. For scRNAseq experiments, 1 million hiPS-CMs and 200 000 ECs per well were plated together on a 12-well plate, and the same amount of hiPS-CMs and hiPS-ECs in monoculture were plated at the same time. After 48 hours of coculture, the cells were used for scRNAseq.

### Processing of cells for single-cell RNA-sequencing

The cells were detached using Accutase (A6964, Sigma) and their concentration was measured. The cells were washed once with PBS containing 0.04% BSA, and then resuspended in PBS with 0.04% BSA to a concentration of 0.79-1.0 x 10^6 cells/ml. The cells were passed through a 35 µm strainer (352235, Corning) and placed on ice until continuation of the 10X Genomics Single Cell Protocol at the Institute of Molecular Medicine Finland (FIMM), where the concentration and viability of cells was calculated with Luna Automated Cell Counter and 4000 cells/sample were processed.

### Single-cell sequencing

Single-cell gene expression profiles were studied using the 10x Genomics Chromium Single Cell 3’RNAseq platform. Data processing and analysis were performed using 10x Genomics Cell Ranger v2.1.1 pipelines. Analyses were performed with Seurat R package version 3.0.1^24^. The detailed methods for single-cell analyses are presented in the Online Data Supplement.

### Single cell sequencing - data presentation

For clarity we present the results in the main figures for the cell line HEL47.2. Only results which were statistically significant in both cell lines are presented, unless otherwise stated. The corresponding results for the cell line HEL24.3 are presented in the Online Data Supplement. Separate replication experiments for flow and coculture were conducted for HEL24.3. To demonstrate that including flow cells in the clustering in HEL47.2 cell line did not affect our results and conclusions, we have repeated the clustering in HEL47.2 also without flow-exposed cells, and present these results in the Online Data Supplement along with HEL24.3 results.

### RNA-sequencing

RNAsamples were sequenced with Illumina NextSeq sequencer (Illumina, San Diego, CA, USA). The detailed methods for RNA Seq analyses are presented in the Online Data Supplement.

## Results

### Characterization of the hiPS-CMs and hiPS-ECs

Immunofluorescence staining of the hiPS-ECs showed strong expression of VE-cadherin and PECAM1 in cell junctions, whereas the hiPS-CMs expressed alpha-actinin and had clear sarcomeric structures (Figure 1A). In addition, the hiPS-ECs were able to take up oxidized LDL (Figure 1A), demonstrating their functionality. Matrigel tube assay showed that the hiPS-ECs were capable of forming tubes in Matrigel similarly to HUVECs (Figure 1C). Interestingly, although the hiPS-EC tubes were not as uniform, they lasted longer than the HUVEC-derived structures. qPCR analysis showed a similar increase in the expression of cardiac genes in the hiPS-CMs during differentiation (Figure I in the Data Supplement). Even though the hiPSC-derived cells do not fully recapitulate the phenotype and function of the respective adult cell type, they provide an excellent tool to model tissue development, and in this study the organotypic development of cardiac ECs and the interaction between ECs and CMs.

**Figure 1.**
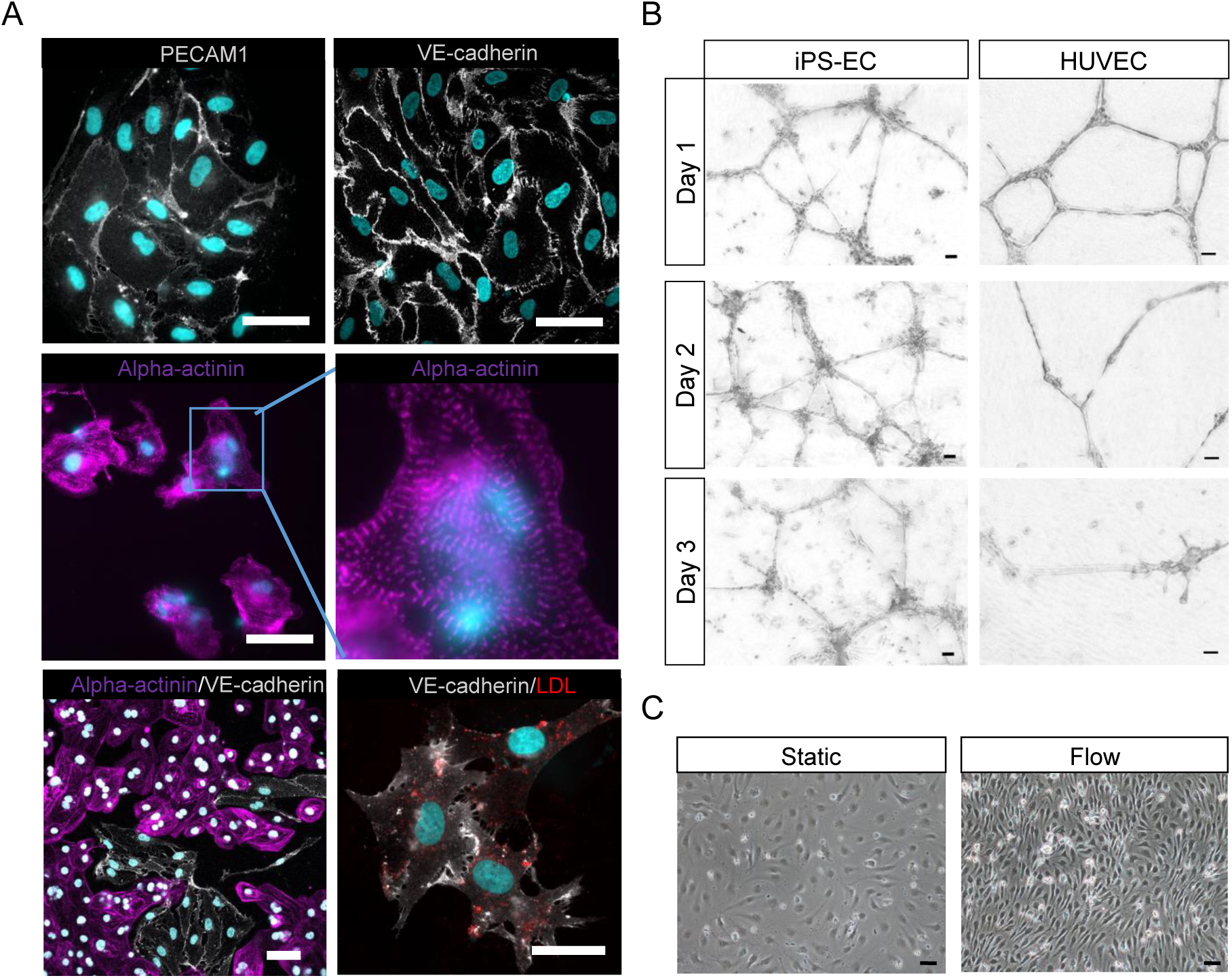
Characterization of iPS-ECs. **A**, PECAM1 and VE-cadherin IF staining of iPS-ECs, alpha-actinin staining of iPS-CMs. The cells are either cultured alone or together in coculture. iPS-ECs take up oxLDL (red). Scale bar 50 µm. **B**, Matrigel tube assay of iPS-ECs and HUVECs (scale bar 50 µm). The cells were plated on Matrigel and imaged every day for three days. **C**, Images of iPS-ECs in static conditions and after 24 hours of flow-induced shear stress. Scale bar 50 μm.

### Identification of single-cell clusters

To analyze the effect of coculture on hiPS-ECs and hiPS-CMs, and the effect of flow-induced shear stress on hiPS-ECs, we performed single-cell RNA sequencing. We compared hiPS-ECs and hiPS-CMs in monoculture to those in coculture, and hiPS-ECs in static monoculture to hiPS-ECs exposed to flow. A total of 14 clusters were identified in the aggregated data of all cell types and treatments combined (Figure 2A through 2E). hiPS-ECs formed separate, distinct clusters based on their growth conditions (Figure 2A and 2B, and Figure IIA and IIB in the Data Supplement) with the exception of proliferating cells, and cells with a high mitochondrial gene content; these clusters contained cells from all conditions. In contrast, mono- and coculture hiPS-CMs clustered together, and two clear hiPS-CM clusters were identified based on their identity (Figure 2A and 2C).

**Figure 2:**
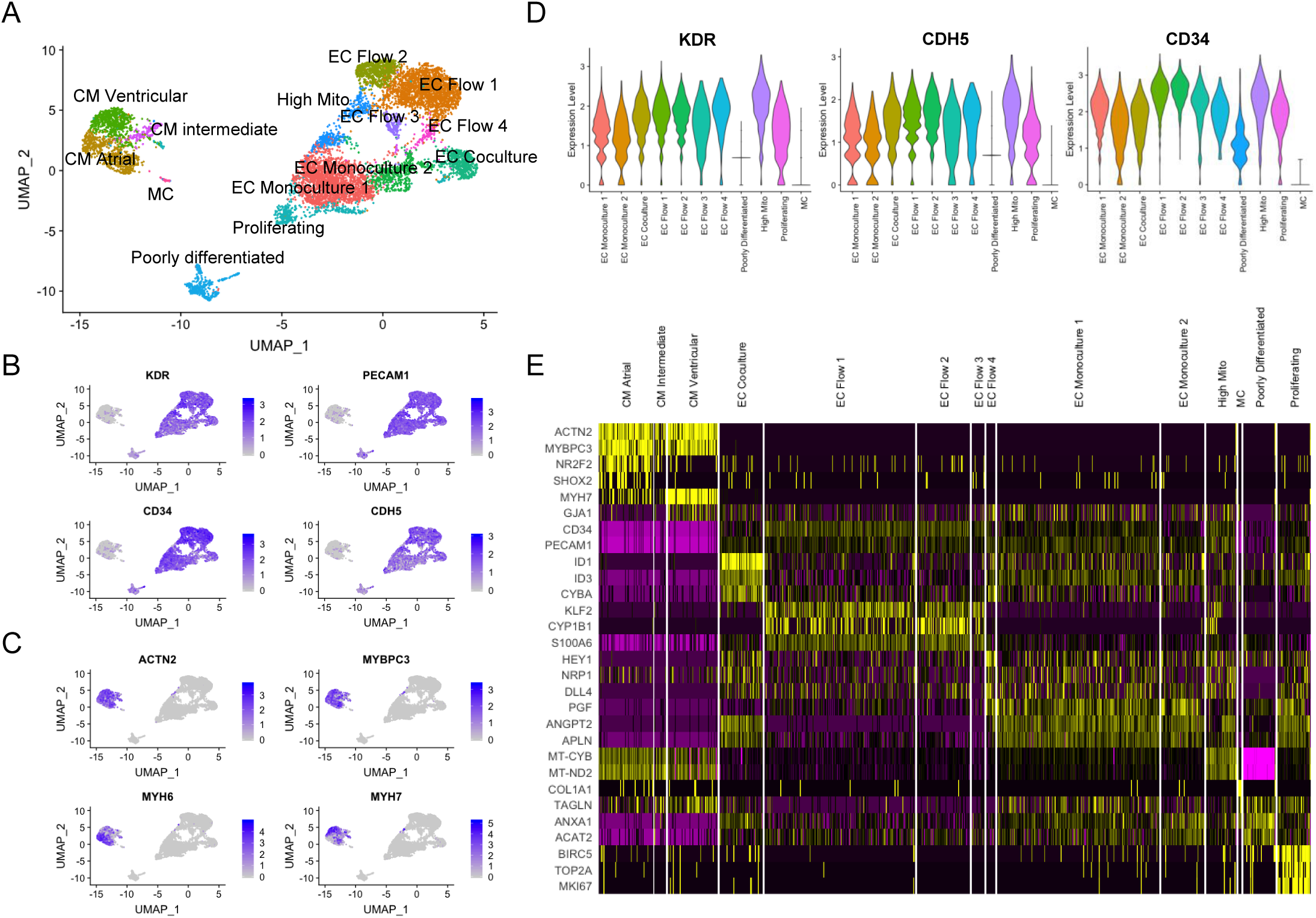
Determining the scRNASeq clusters (HEL47.2). **A**, UMAP (Uniform Manifold Approximation and Projection) plot of single cell sequencing. **B**, Feature plot of EC gene expression. **C**, Feature plot of CM gene expression. **D**, Violin plots of expression of general EC markers in clusters including iPS-ECs. **E**, Heatmap on the expression of representative genes clusterwise.

The hiPS-ECs had high expression of the endothelial cell markers *PECAM1, KDR, CD34,* and *CDH5* (Figure 2B and 2D). Out of the 14 clusters, seven were identifiend with only hiPS-ECs (Figure IIB in the Data Supplement). Two of the hiPS-EC clusters consisted mainly of cells from static monoculture (EC Monoculture 1-2), one hiPS-EC cluster consisted mainly of coculture cells (EC Coculture), and four distinct hiPS-EC were comprised almost solely of cells exposed to flow (EC Flow 1-4). The EC Monoculture 2 cluster had a small number of cells from flow and coculture, probably representing cells that did not respond to stimuli.

The hiPS-CMs formed two distinct clusters, of which one had an atrial (CM Atrial) and the other a ventricular (CM Ventricular) gene expression profile (Figure 3A-C). The ventricular cluster had higher expression of *MYL2, GJA1, HEY2*, and *IRX4* ^25,26^, whereas the atrial cluster had higher expression of *NR2F2, KCNJ3, CACNA1D*, and *SHOX2* ^*25,27*^ Optogenetic analysis of the hiPS-CM confirmed the presence of both atrial and ventricular cells with respective action potential features (Figure 3D).

**Figure 3:**
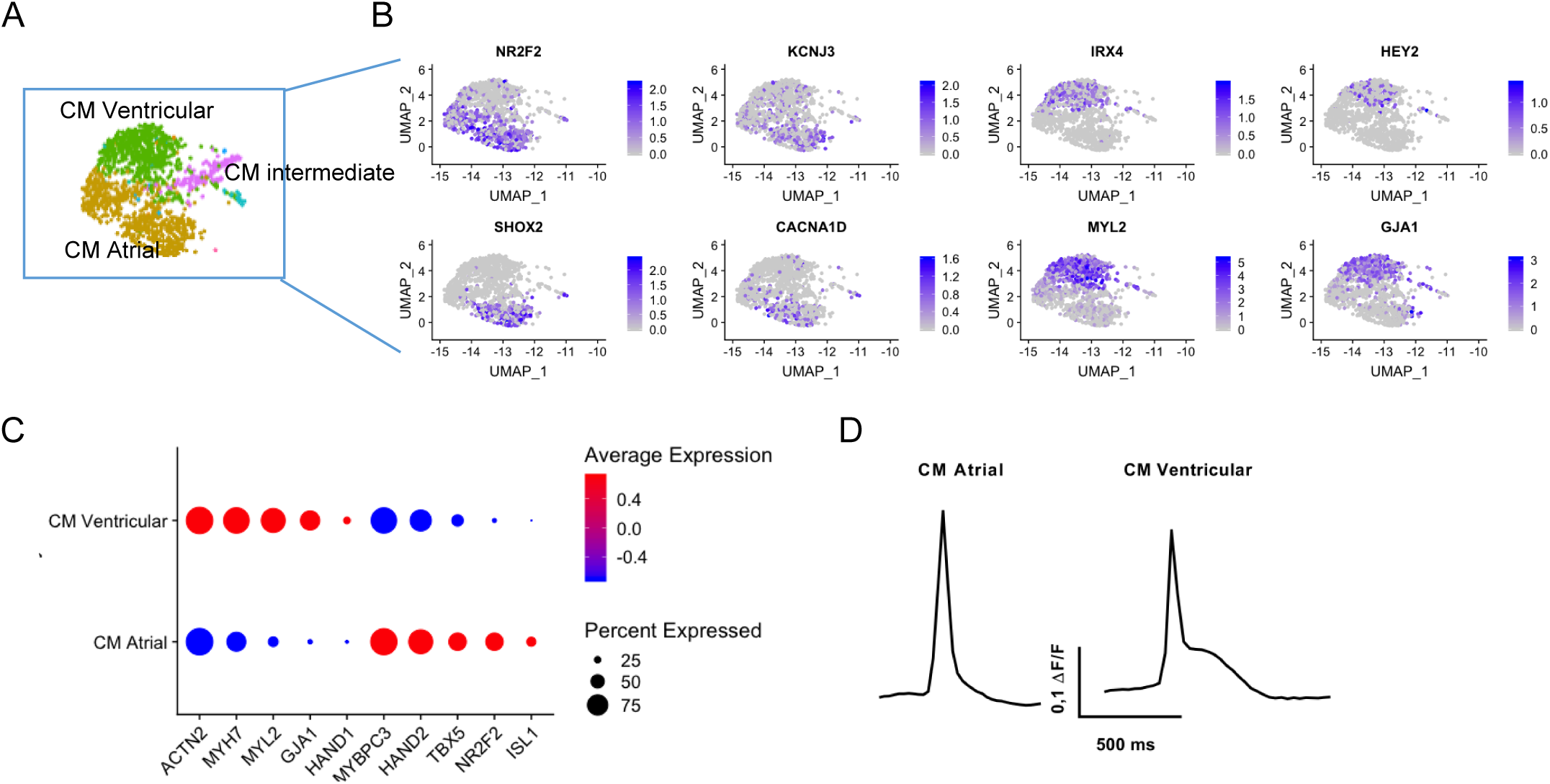
Characterization of the CM clusters of scRNAseq. **A**, UMAP (Uniform Manifold Approximation and Projection) plot of single cell sequencing of CMs. **B**, Feature plot of CM gene expression**. C**, dot plot on the expression of atrial and ventricular CM-genes in the iPS-CM clusters. **D**, Representative action potential morphologies of atrial and ventricular iPS-CMs.

Five clusters had clearly mixed cell populations. Proliferating cells (Proliferating) with high expression of the cell cycle genes *MKI67, TOP2A*, and *BIRC5* included cells from mainly static hiPS-ECs and coculture hiPS-ECs (Figure IIB in the Data Supplement) reflecting the flow-induced inhibition on EC proliferation. Cells from all groups were represented in the cluster of cells containing a high mitochondrial gene content (High Mito) (Figure IIB in the Data Supplement). The majority of these cells were static or flow hiPS-ECs, and might represent an unhealthy population, which would have been excluded if the mitochondrial gene content limit was set lower. In addition, there was a distinct cluster of cells with low EC gene expression, which we named as poorly differentiated, mainly consisting of static hiPS-ECs (Figure 2A and Figure IIB in the Data Supplement). Finally, there was a small cluster between atrial and ventricular hiPS-CMs (CM Intermediate), and a small cluster close to hiPS-CMs (MC - Mesenchymal cells), which included cells mainly from the hiPS-EC samples and expressed a mesenchymal cell phenotype with high expression of *TAGLN*, and *ACTA2*.

Highly similar clustering was found in the HEL24.3 cell line and when the HEL47.2 cell data was clustered without flow-exposed cells (Figure III, and Figure IVA, and IVB in the Data Supplement). The CM Atrial cluster, however, was a lot smaller in the HEL24.3 cell line, and there was a larger cluster of CMs with a gene expression profile between atrial and ventricular CMs (Figure IVC and IVD in the Data Supplement), which potentially indicates a lower degree of maturation.

### The effect of coculture on hiPS-EC and hiPS-CM transcriptomes

The differential gene expression analysis revealed a more pronounced effect of coculture in the hiPS-EC transcriptome compared to that of hiPS-CM. As compared with hiPS-ECs grown in monoculture the expression of 130 genes were increased and 68 genes repressed in both cell lines, HEL47.2 (318 up, 155 down) and HEL24.3 (321 up, 187 down) (Table II in the Data Supplement).

A gene ontology classification analysis of the 130 genes upregulated in hiPS-ECs in both cell lines revealed activation of pathways associated with cardiovascular development, cell signaling, angiogenesis, and cell differentiation (Figure 4A). More specifically, upregulation of general EC genes (Figure 2D), and, interestingly, cardiac EC-specific genes^28^ (Figure 4B), as well as several genes associated with heart development^29–34^ (Figure 4C), and angiogenesis^35–37^ (Figure 4D) were observed in hiPS-ECs cultured together with hiPS-CMs. The respective data for the HEL24.3 cell line are presented in Figure V in the Data Supplement.

**Figure 4:**
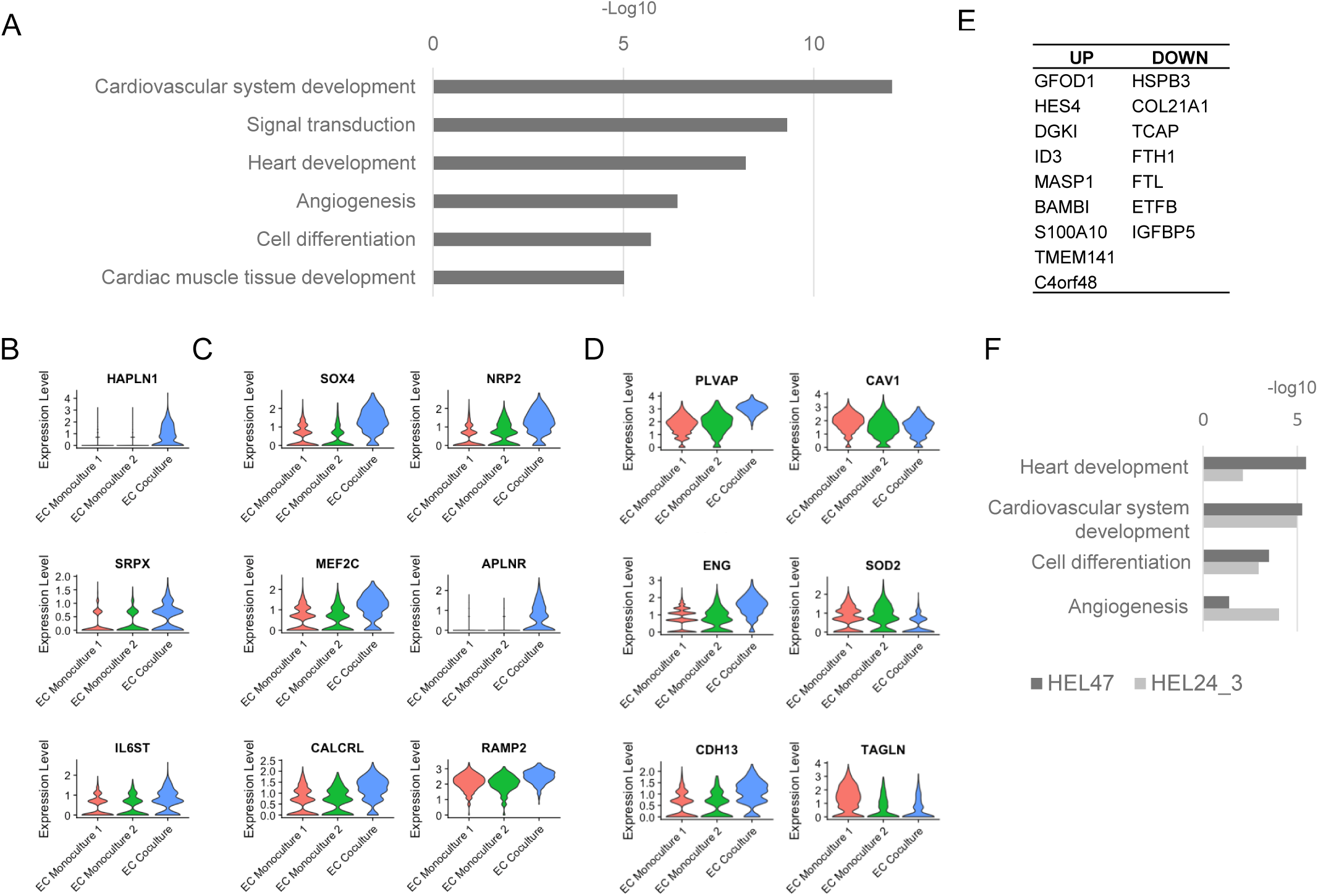
The effect of coculture in iPS-EC and iPS-CM gene expression (HEL47.2). **A**, GO analysis of genes that were upregulated in iPS-ECs in both cell lines (HEL47 and HEL24_3) in coculture. Coculture led to upregulation of **B**, cardiac specific EC-genes, **C**, genes associated with heart development, and **D**, genes associated with regulation of angiogenesis. **E**,Genes that were similarily up- and downregulated in HEL47.2 and HEL24.3 in cocoulture iPS-CMs. **F**, GO analysis of genes that were upregulated HEL47.2 and HEL24.3 iPS-CMs in coculture.

Coculture resulted in upregulation of 39 and 40 genes, and downregulation of 30 and 24 genes in HEL47.2 and HEL24.3 hiPS-CMs, respectively, compared to monoculture hiPS-CMs (Table III in the Data Supplement). Of these, 9 were upregulated and 7 downregulated in both cell lines (Figure 4E). Augmented expression of genes involved in heart and cardiovascular system development were observed in the coculture hiPS-CMs (Figure 4F).

### Coculture reprogrammed BMP-signaling pathway in hiPS-CMs and hiPS-ECs

Cells grown in coculture exhibited changes in the BMP-signaling pathway, which is an important regulator of cardiac development. The ligand *BMP4* was markedly increased in coculture hiPS-ECs. In hiPS-CMs, the expression of *BMP2* was restricted to atrial cells and it increased significantly in HEL47.2 coculture, whereas the expression levels of *BMP5*, and *BMP7* remained constant (Figure 5A). The hiPS-ECs and hiPS-CMs expressed several BMP-pathway receptors, and their expression was not significantly affected by the presence of the other cell type (Figure 5B). Interestingly, the expression of the decoy receptor *BAMBI* increased in both cell types. Of the BMP-signaling effectors, the expression of *SMAD1* was lower in coculture hiPS-ECs, whereas the expression levels of other SMADs remained unchanged (Figure 5C). The expression of BMP targets *ID1, ID2*, and *ID3* were all significantly increased in coculture hiPS-ECs and the expression of *ID3* was higher in the coculture hiPS-CMs (Figure 5D). The respective violin plots for the HEL24.3 cell line are presented in Figure VI in the Data Supplement.

**Figure 5:**
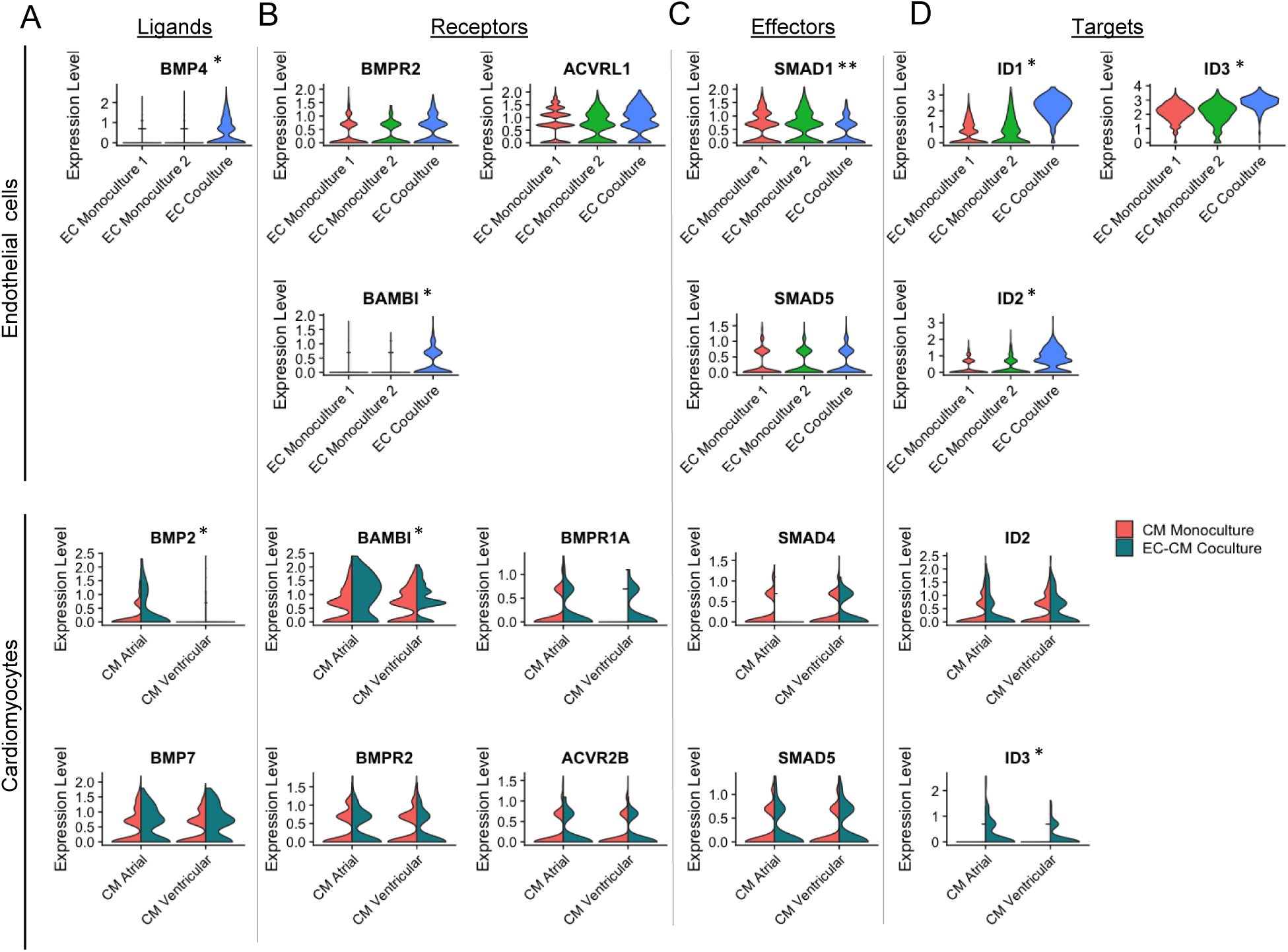
The expression of BMP-pathway genes in iPS-ECs and iPS-CMs in monoculture and coculture (HEL47.2). Violin plots of the expression of BMP **A,** ligands, **B,** receptors, **C,** effectors, and **D,** targets. * = Siginificantly higher expression in coculture vs. monoculture cells (adjusted p<0.05). **= Siginificantly lower expression in coculture vs. monoculture cells (adjusted p<0.05).

Since the BMP pathway was found to be affected in coculture, we checked the expression levels of other BMP-pathway ligands, receptors, effectors and targets. Most of their levels were low or undetectable (Table IV in the Data Supplement).

### Coculture increases the expression of NOTCH-pathway genes but not ERBB-pathway

During cardiac development, the interaction between ECs and CMs has been shown to be especially important for myocardial trabeculation, where NOTCH-ERBB pathways have been demonstrated to be important mediators of the crosstalk between endocardium and myocardium. Thus, we next examined the effect of coculture on the expression of the NOTCH and ERBB pathway mediators. Significant increase in the NOTCH-ligand *JAG1* was observed in HEL47.2 coculture hiPS-CMs, whereas in the hiPS-ECs there was a trend for increased expression of *JAG1* in both cell lines (Figure VII in the Data Supplement). The hiPS-ECs expressed NOTCH receptors 1 and 4 in both conditions. Of NOTCH-pathway targets, *HES4* expression levels were higher in the coculture hiPS-CMs in both cell lines compared to monoculture cells. In addition, higher expression levels of NOTCH target *HEY2* were observed in coculture hiPS-ECs in both cell lines (Figure VII in the Data Supplement). Higher expression of the target *HES1* was observed in HEL47.2 coculture hiPS-ECs and *HES4* in HEL24.3 coculture hiPS-ECs.

*ERBB2* and *ERBB4* were expressed in hiPS-CMs, and the expression levels were similar in both atrial and ventricular clusters. *ERBB3* was also expressed at low levels in hiPS-CMs. Of the ERBB pathway mediators, only *HB-EGF* was expressed at detectable levels in the hiPS-ECs. We did not observe significant changes in the ERBB pathway gene expression in coculture cells, either hiPS-CMs or hiPS-ECs (data not shown).

### Exposure to flow results in both arterial- and venous-like hiPS-ECs

As ECs are constantly exposed to flow *in vivo*, we studied the transcriptomic changes in hiPS-ECs exposed to flow as compared to static monoculture hiPS-ECs (Figure 6A). This was done by scRNASeq in the HEL47.2 and HEL24.3 cell lines. In addition, the flow-experiment was replicated in four hiPS-EC-lines (HEL47.2, HEL24.3, HEL46.11, and K1), and analysed with bulk RNASeq. Clustering of the HEL24.3 EC flow and EC monoculture cells is presented in Figure VIIIA in the Data Supplement.

**Figure 6:**
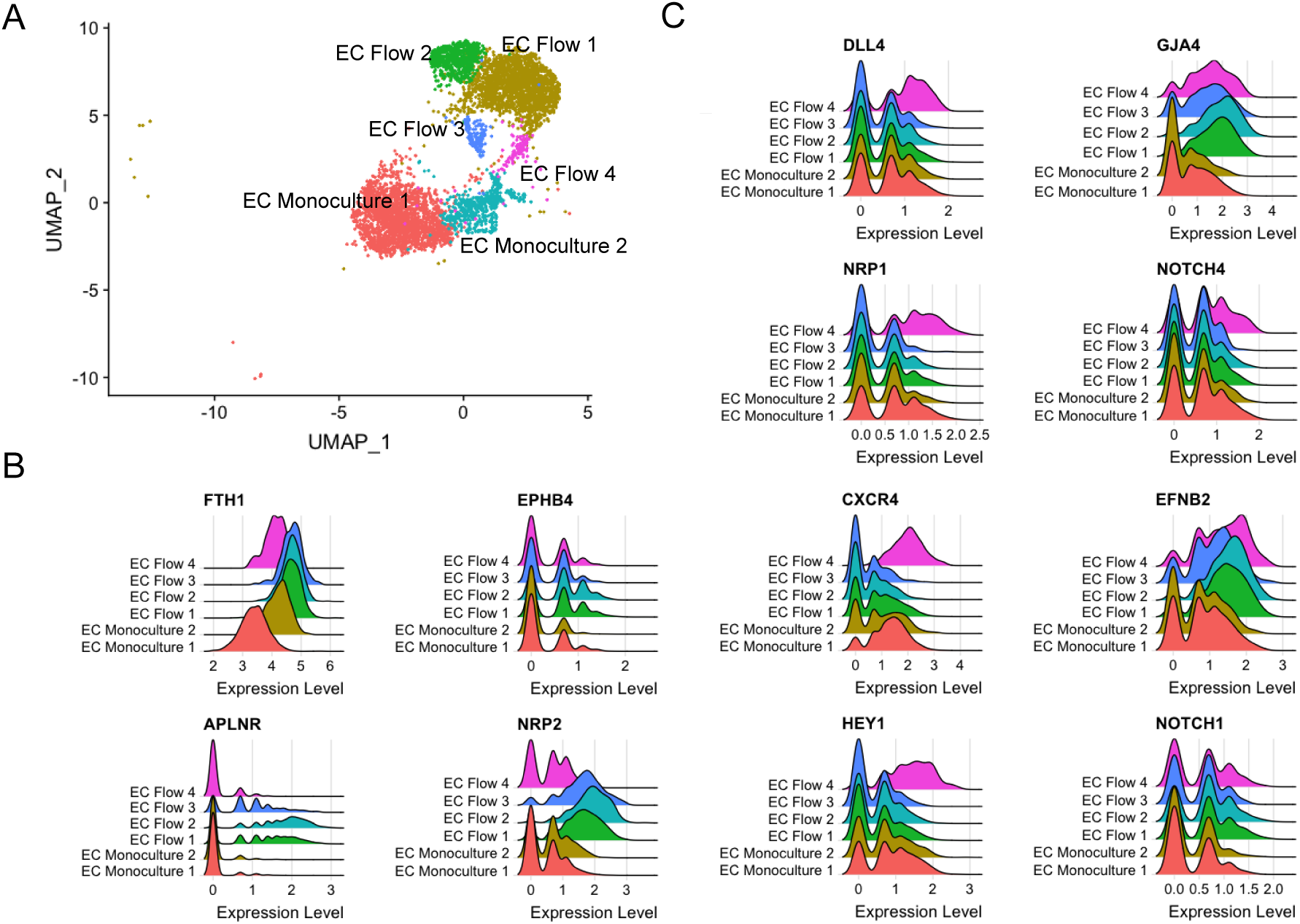
The expression of arterial and venous genes in static monoculture iPS-EC and monoculture iPS-ECs exposed to flow. **A,** UMAP of the custers included in the analysis. Ridge plots of the expression of **B** venous genes and **C,** arterial genes.

Flow resulted in upregulation of 174 genes, and downregulation of 105 genes in the scRNASeq (common genes in both cell lines). When the bulk RNASeq data was analyzed together, in total 101 genes were upregulated and 56 genes were downregulated in all data sets, forming a core of the flow-induced genes (Table V in the Data Supplement)

According to the scRNASeq analysis, flow increased the expression of characteristic EC markers (Figure 2D), and both arterial and venous EC markers. HEL47.2 EC Flow clusters 1-3 had higher expression of venous genes (Figure 6B), while the cluster EC Flow 4 had a higher expression of arterial genes (Figure 6C). Similar clustering was found in the HEL24.3 hiPS-ECs with one arterial-like cluster (Figure VIIIB and VIIIC in the Data Supplement). In the bulk RNASeq data, where the effects of flow were studied as single population, significant expression differences were found for the venous markers *NRP2, FTH1*, and *EPHB4*, and the arterial marker *NOTCH1* (Table V in the Data Supplement). Interestingly, except for the scRNASeq clusters EC Flow 4 in the HEL47.2 cell line and EC Flow 5 in the HEL24.3 cell line, which showed enhanced arterial phenotype, the expression of arterial genes were not significantly changed in the flow hiPS-Ecs compared to static hiPS-ECs. This demonstrates the power of scRNAseq, as the arterial cluster was relatively small, and these changes cannot be detected from the bulk RNAseq data. The top 25 up- and downregulated genes in flow hiPS-ECs in the bulk RNASeq analysis are presented in Figure IX in the Data Supplement.

Shear stress activated pathways associated with cardiovascular development, angiogenesis, and cell signalling (Figure 7A). We then looked at genes that were significantly up- and downregulated in both scRNASeq cell lines and the bulk RNASeq. The expression of known flow-induced genes *KLF2, KLF4*, and *eNOS* (*NOS3*), which promote vascular tone^38^, were highly upregulated in flow (Figure 7B). In addition, the expression of the stress response markers *HMOX1*^*39*^, and *NQO1*^*40*^, the anti-atherogenic genes *CYP1A1*, and *CYP1B1*^*41*^, and genes that promote vascular homeostasis such as *SLC9A3R2* (*NHERF2*)^42^ and *TIMP3*^*43*^ were upregulated in flow (Figure 7B). In contrast, shear stress downregulated the vascular tone regulator *EDN1*^*44*^ in all flow clusters (Figure 7C). Flow promoted quiescence in endothelial cells and this was consistent with the finding of several angiogenesis genes such as *APLN*^*45*^, *ANGPT2, CITED2*^*46*^, DDAH1^47^, and *THBS1*^*48*^ being downregulated by flow (Figure 7C). The arterial-type FlowEC 4 cluster seemed to have attenuated response to flow, based on the lower expression of flow-induced genes compared to the other three flow clusters, however they still clearly differed from the static cells. The respective violin plots for the HEL24.3 cell line are presented in Figure X in the Data Supplement.

**Figure 7:**
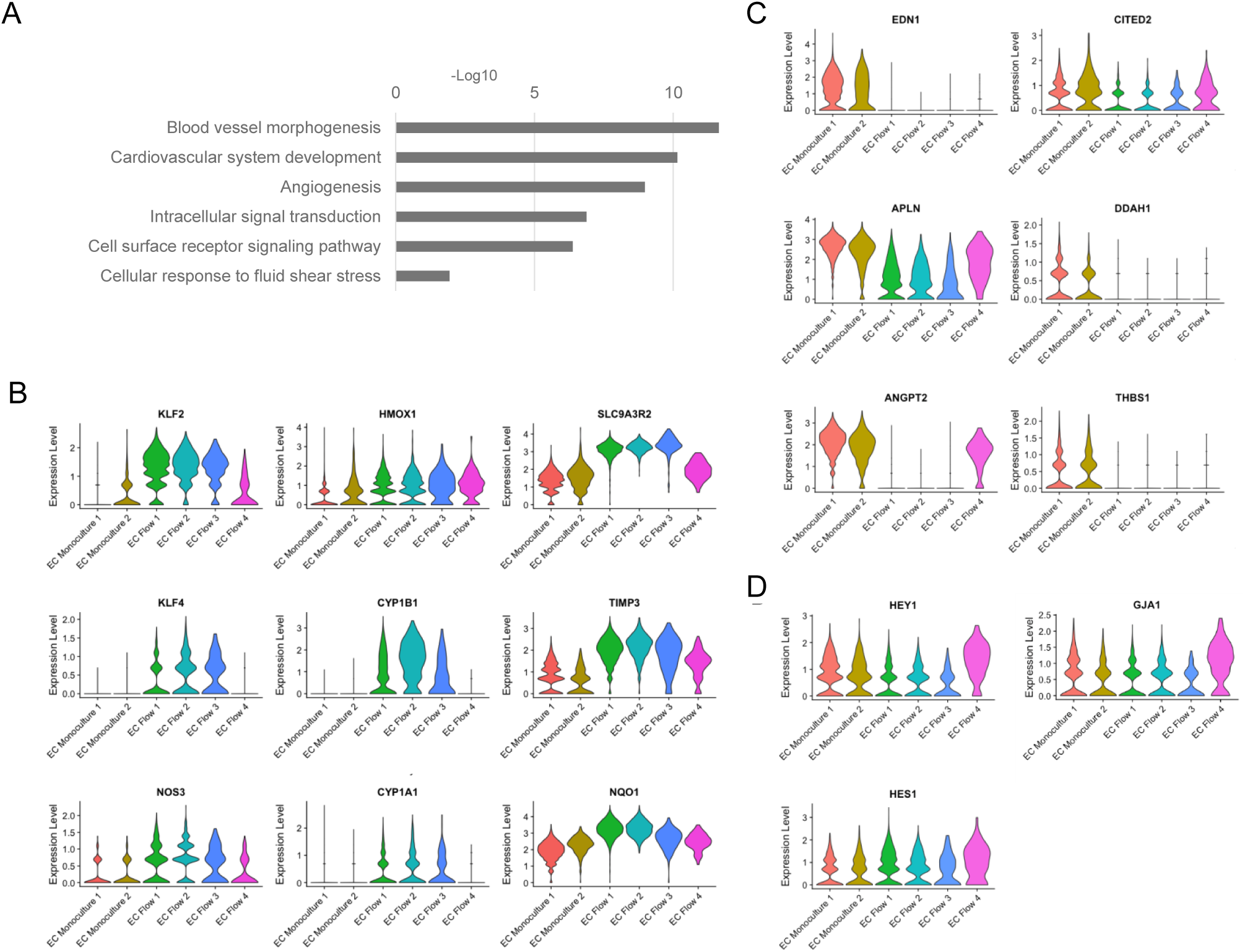
Shear stress induced significant changes in the iPS-EC transcriptome. **A,** GO analysis of genes that were upregulated in iPS-ECs in both cell lines (HEL47 and HEL24_3). In both scRNASeq cell lines and bulk RNASeq, shear stress led to **B,** upregulation of known flow-induced genes, antiatherogenic genes and genes that promote vascular homeostasis, and **C,** downregulation of vascular tone regulators and angiogenesis genes. **D,** Increased expression of NOTCH-targets was observed in the flow-exposed arterial-like iPS-Ecs (EC Flow 4).

### Flow induces NOTCH-signaling in hiPS-ECs

Shear stress is a known NOTCH-pathway activator, and it has been shown to increase *NOTCH1* gene expression in endothelial cells. Compared to static hiPS-ECs, altered expression of several NOTCH pathway genes were observed in the hiPS-ECs exposed to flow. Interestingly, the effects were different in the venous-like (EC Flow 1-3) and arterial-like (EC Flow 4) clusters. The notch-ligand *DLL4* was upregulated in the arterial-like hiPS-ECs in flow, whereas the expression level was lower in the venous-like hiPS-ECs compared to hiPS-ECs in static conditions (Figure 6A). The expression levels of *NOTCH1* and *NOTCH4* receptors were upregulated in all flow hiPS-EC clusters, but more so in the arterial-like cluster (Figure 6A). Likewise the expression of the NOTCH-targets *HEY1, HES1*, and *GJA1* was higher in the arterial-like FlowEC 4 (Figure 7D).

### Common effects on EC transcriptome by coculture with CM and exposure to flow

A total of 32 same genes were upregulated and 28 downregulated in both coculture and flow exposed hiPS-ECs in both cell lines (Figure 8A and 8B and Table VI in the Data Supplement). The most significantly induced genes by both coculture and flow included genes that are implicated in vascular homeostasis, and heart development (Figure 8C). *PRCP, APLNR, PLVAP, HAPLN1,* and *CD59* are important in vascular health and regeneration^33,34,36,49,50^. *PLVAP* is also highly expressed in adult endocardium^36^. *HAPLN1 (CRLT1)* has been shown to be expressed in endocardium during heart development, regulating cardiac cushion formation, and is along with *APLNR,* and *ADAMTS9* essential during cardiac development^29,33,51^. Moreover, flow and coculture both induced downregulation of atherogenic genes, such as *CAV1*^*35*^, and oxidative stress induced genes such as *PGF*^*52,53*^and *SOD2* ^37^ (Figure 8D).

**Figure 8:**
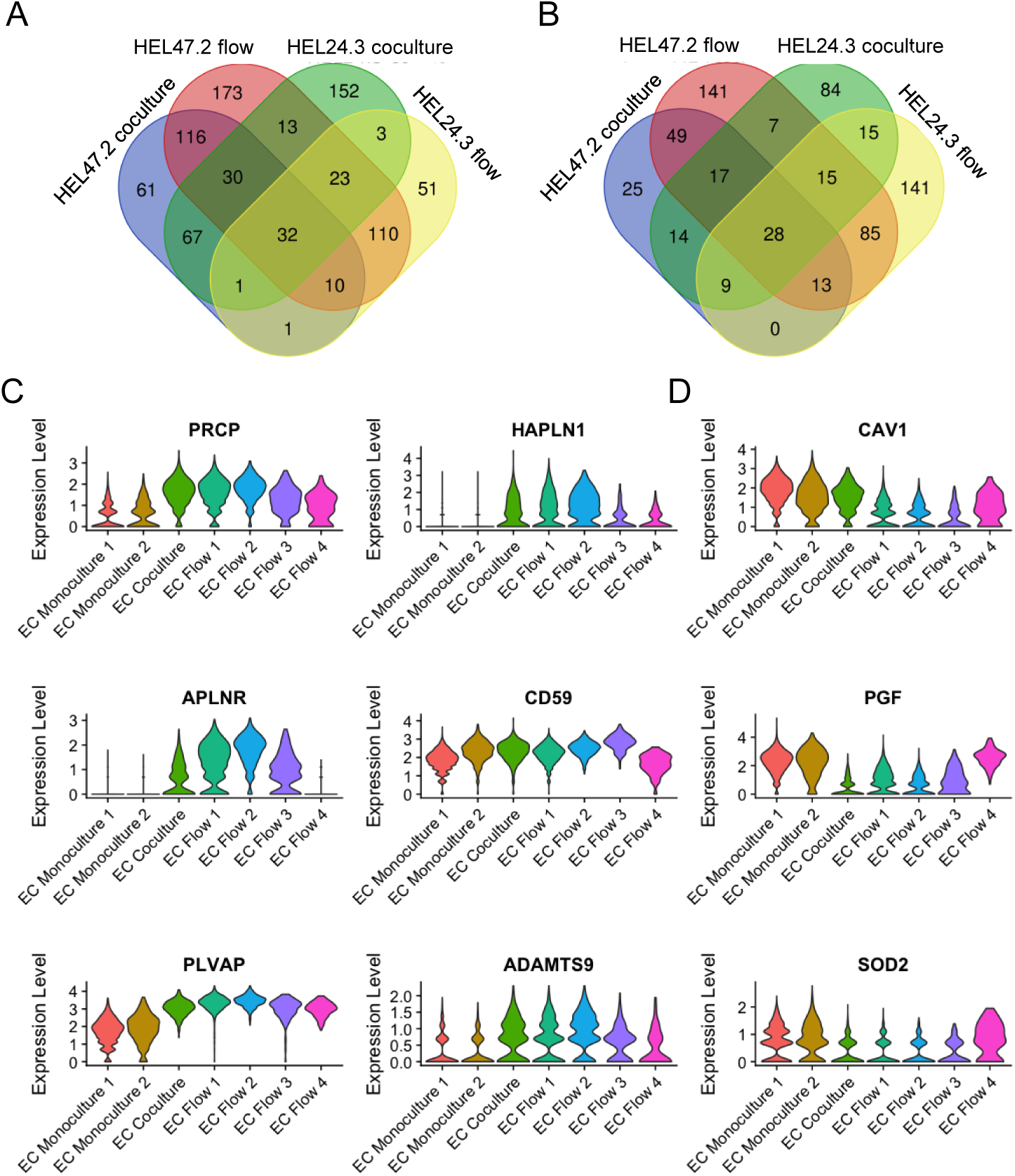
Common effects of flow and cocolture on iPS-EC transcriptome. **A,** Venn diagram of no. of upregulated genes in flow and coculture in both cell lines **B,** Venn diagram of no. of downregulated genes in flow and coculture in both cell lines **C,** Similarily upregulated genes included those implicated in vascular homeostasis and heart development. **D,** Common downregulated genes associated with vascular tone regulation and angiogenesis.

## Discussion

Patient derived hiPS-ECs are increasingly used in disease modeling to study specific disease related phenotypes and genotype-phenotype correlations. However, hiPS-ECs do not gain full maturity and identity compared to ECs in vivo, where ECs are exposed to mechanical forces of blood flow such as shear stress and pulsatile pressure, and paracrine signals from neighboring parenchymal cells. In addition, the efficiency of differentiation as well as maturity of cells vary depending on the protocol used. Here we demonstrate that differentiating hiPSCs to hiPS-ECs results in a population of cells with cobble-stone morphology, and high expression of EC molecular markers. In addition, these cells are able to form vascular tube structures in 3D and take up oxidized LDL. We show that the hiPS-ECs have high plasticity as they adapt to flow and coculture with hiPS-CMs by entering quiescence and further increasing their EC marker expression. Interestingly, the expression of both arterial and venous genes increased in flow, and clustering revealed flow-exposed subpopulations with transcriptomic profiles towards either a more venous or arterial identity. The hiPS-EC transcriptomic landscape was also strongly affected by coculture with hiPS-CMs, resembling organotypic adaptation to the requirements of the parenchymal cells. This was reflected by augmented expression of genes important during cardiac development, and genes that are specifically expressed in cardiac ECs. Our results highlight the importance of single-cell RNA sequencing to distinguish different phenotypes of hiPSC-derived ECs and CMs and the potential of coculture in disease modeling.

Nearly all hiPS-ECs highly expressed established endothelial markers *PECAM1, CDH5,* and *KDR*. The expression levels of the endothelial progenitor marker *CD34* were high, which is in accordance with previous studies demonstrating an immature nature of these cells, potentially resulting in plasticity to further develop to, or even transfer between arterial and venous phenotypes ^15,16^. In contrast, the expression levels of the lymphatic EC markers *PROX1* and *PDPN* were very low, indicating that there were no lymphatic ECs among the hiPS-ECs. A small number of cells expressed the smooth muscle cell marker *ACTA2*, and these cells clustered far from other hiPS-ECs. Monoculture hiPS-ECs under static conditions expressed more arterial than venous markers, which is consistent with previous reports ^54,55^. However, the cells did not cluster according to arterial or venous phenotype in static monoculture.

Single-cell RNA sequencing of flow exposed hiPS-ECs revealed increased expression of several arterial and venous genes, with higher induction seen in venous markers, which is in accordance with previous studies ^56^. The expression pattern was replicated in the bulk RNASeq data for the venous markers *NRP2, FTH1,* and *EPHB4,* and the arterial marker *NOTCH1*. Importantly, in the single-cell sequencing data, several hiPS-EC-subclusters were identified under flow, of which one was clearly arterial-like and the others venous-like, underlining the responsiveness of the cells to external stimuli and also the heterogeneity of the phenotypic change. As expected, flow activated known vascular tone regulators, such as *KLF2* and *KLF4*^*38*^, and shear responsive anti-atherogenic genes, such as *CYP1A1* and *CYP1B1* ^41^. Interestingly, it seems that those cells, which had smaller responses to flow, e.g. attenuated upregulation of *KLF2* and *CYP1B1*, became arterial-like cells.

Cardiomyocytes formed two main clusters, which according to their gene expression profiles could be defined as atrial and ventricular. The atrial cluster expressed the transcription factors *SHOX2*, and *NR2F2*, and the atrial ion channels *KCNJ3*, and *CACNA1D*^*25,27,57*^. In the adult human heart, all of these four genes are mainly expressed in the human atrial appendages (GTEx V8, gtexportal.org/home/). *MYH6* and *TBX5* levels were higher in atrial cells, and *MYH7, MYL2*, and *GJA1* in ventricular cells^25,26^, corresponding with their expression in the adult human heart (GTEx V8). HAND proteins HAND1 and HAND2 are closely related transcription factors that play critical roles during cardiogenesis, and contribute to the maturation of several substructures within the developing heart, including the cardiomyocytes of the ventricles, myocardial and cardiac neural crest contributions to the OFT, and epicardium ^58^. *HAND2* was expressed in both atrial and ventricular hiPS-CMs, as found in the adult human heart (GTEx V8). Consistently with the human adult heart, *HAND1* expression was almost solely limited to ventricular-like hiPS-CMs. The transcriptomic data was recapitulated by optogenetic analysis of IPC-CMs electrophysiology, which demonstrated the presence of both atrial and ventricular-types of action potentials within the cardiomyocyte populations.

Endothelial cells and cardiomyocytes communicate with each other by cardiokine and angiocrine signaling, and this crosstalk has an essential role in cardiac development, growth, and homeostasis. Significant transcriptomic shifts were observed in both cell types in coculture, however, the differences were more pronounced in hiPS-ECs. This demonstrates the plasticity of ECs also *in vivo* to adapt to their environment by acquiring organotypic features in different tissues. It also likely reflects the developmental state of the hiPS-ECs with high plasticity towards paracrine signals. Interestingly, although the hiPS-CMs are considered to resemble embryonic cardiomyocytes, they had a markedly smaller response to coculture than the hiPS-ECs.

BMP/TGF-β- and NOTCH-pathways are among the most studied in cardiac morphogenesis. We observed changes in the expression of several BMP/TGF-β and NOTCH pathway ligands, receptors, targets, and effectors in both cell types in coculture. While continuous BMP-signaling is required during early cardiogenesis, later atrial and ventricular cardiomyocytes require BMP signaling at different stages ^59^. Interestingly, we observed differences in atrial and ventricular hiPS-CMs in BMP signaling. For example, BMP2 was mainly expressed in atrial-type hiPS-CMs. In coculture, increased expression of the decoy receptor BAMBI occurred mainly in atrial-type cardiomyocytes, and the expression of SMAD4 diminished in ventricular-type cardiomyocytes. This demonstrates the heterogeneity of these cells, and possibly indicates a cell-type specific regulatory effect of hiPS-ECs on hiPS-CM BMP-signaling. Markedly increased levels of several ID genes in both hiPS-ECs and hiPS-CMs in coculture further underline the strong effect these cells have on each other.

NOTCH-signaling is a typical example of the role of EC-CM crosstalk during cardiac development. NOTCH-signaling has a role in the patterning of the early embryonic endocardium to valve and chamber formation, and later regulating the outflow tract and valve morphogenesis^60^ and ventricular trabeculae compaction^61^. Increased expression of several NOTCH-pathway ligands and targets in both cell types demonstrate the activation of this pathway in coculture. The expression of Notch ligands *JAG1* and *DLL4* in hiPS-ECs, and the higher expression of *JAG1* in hiPS-CMs in coculture resembled that of early phases of cardiogenesis in mice^62^. Interestingly, the expression of *JAG1*-ligand was higher in the atrial-like hiPS-CMs, which is consistent with the gene expression pattern in murine embryonic hearts^63^.

Augmented expression of several other genes associated with heart development^30–34,50,51^ such as *SOX4, MEFC2, RAMP2, CALCRL*, and *APLNR* and organotypic cardiac endothelial cell genes ^28^, such as *HAPLN1, SPRX1* and *IL6ST*, was observed in coculture hiPS-ECs. Many of the coculture induced genes are also known to contribute to vascular development and angiogenesis, likely reflecting their response to angiogenic factors produced by cardiomyocytes.

Both shear stress and the presence of hiPS-CMs led to upregulation of genes associated with vascular homeostasis and quiescence in hiPS-ECs. Of these, PRCP has a role in vascular growth and repair after injuries ^49^, and PLVAP (Plasmalemma vesicle-associated protein) is an endothelial cell-specific protein that forms the stomatal and fenestral diaphragms of blood vessels and regulates basal permeability, leukocyte migration and angiogenesis ^36^, and it is upregulated in pathogenic angiogenesis ^64^. Repression of genes associated with hypoxia and other pathologic cardiac conditions were also seen in both flow and coculture-exposed hiPS-ECs ^37,52,53^.

As expected, the responses to stimulation were not fully identical in the studied cell lines. It has been previously reported that the results can vary even within a cell line between two differentiations. However, most of the major effects observed in our study replicated in both cell lines. The EC clustering was similar in both cell lines, as the flow cells and coculture cells clearly formed their own clusters separate from the static monoculture ECs. In addition, both cell lines had a cluster of hiPS-ECs consisting mainly of static monoculture hiPS-ECs that had lower expression of EC markers, and thus defined as poorly differentiated, and a proliferating hiPS-EC cluster. An interesting observation in the hiPS-CMs was that the Atrial CM cluster was rather small in the HEL24.3 cell line compared to HEL47.2 cell line, and the HEL24.3 cell line had a larger cluster between the atrial- and ventricular-like CM clusters that resembled more ventricular cells, but also had some atrial gene expression. One possible explanation could be the age difference between the hiPS-CMs, as the HEL24.3 were younger at the time of the experiment, thus, the intermediate cluster could represent a more naive cluster. It could be plausible that the differentiation to atrial and ventricular cells occur when more time is given for the cells to mature.

The disadvantages of using hiPSC-derived ECs compared to primary ECs or cell lines is the immature nature of the cells, their tendency for transdifferentiation, the low proliferation capacity over passaging and the costs to produce them ^65^. This is why the hiPS-ECs were used immediately after differentiation and sorting for the experiments. Despite these limitations, hiPSCs are superior in disease modelling when studying the interactions between cell types, as all the cell types can be derived from the same patient carrying the same genetic variants. Thus, there is a need for the development of better hiPS-EC models.

Our study sheds light on the interactions and transcriptomic changes induced by endothelial cell-cardiomyocyte crosstalk, which is an important regulator of cardiac development and adult cardiac homeostasis in health and disease. Human hiPSC-derived endothelial cells and cardiomyocytes provide an excellent translational model to study organotypic development of cardiac vasculature and the EC-CM interactions. The transcriptomic changes observed in the coculture cells after 48 h demonstrate the importance and magnitude of this signaling. Moreover, the extent of phenotypic change in the hiPS-ECs suggests that striving towards more physiological conditions in cell culture could provide us with more accurate disease models. Finally, the observation of atrial and ventricular hiPS-CM subclusters and arterial and venous hiPS-EC subclusters in the flow-exposed ECs demonstrate the value of using scRNAseq, as specific cell types clearly respond differently to the stimuli.

## Acknowledgments

We thank Ilse Paetau for her help in the cell culture and administrative support. We thank Professor Timo Otonkoski and Docent Ras Trokovic at Biomedicum Stem Cell Center for providing us with three control hiPS-cell lines and Professor Anu Suomalainen-Wartiovaara for the gift of the hiPSC line K1. We also thank Kristiina Dahl and Veera Verkasalo for assistance in the optogenetics experiment. We are grateful to Maija Atuegwu and Docent Katariina Öörni for providing us with oxidized-LDL and help with the LDL uptake assay.

## Sources of funding

Finnish Medical Foundation (EH), Finnish Foundation for Pediatric Research (EH), Finnish Foundation for Cardiovascular Research (EH, ET, RK), University of Helsinki (EH), Academy of Finland (grant 297245, RK), Jenny and Antti Wihuri Foundation, Sigrid Jusélius Foundation (RK), Finnish Cultural Foundation (ET, RK), Instrumentarium Science Foundation (ET)

## Disclosures

None

## Supplemental Materials

Expanded Materials & Methods

Data Supplement Figures I-X

Data supplement Tables I-VII

## Notes

### Competing Interest Statement

The authors have declared no competing interest.

## References

1. le Noble F, Moyon D, Pardanaud L, Yuan L, Djonov V, Matthijsen R, Bréant C, Fleury V, Eichmann A. Flow regulates arterial-venous differentiation in the chick embryo yolk sac. Development. 2004;131:361–375.

2. Red-Horse K, Crawford Y, Shojaei F, Ferrara N. Endothelium-microenvironment interactions in the developing embryo and in the adult. Dev Cell. 2007;12:181–194.

3. Atkins GB, Jain MK, Hamik A. Endothelial differentiation: molecular mechanisms of specification and heterogeneity. Arterioscler Thromb Vasc Biol. 2011;31:1476–1484.

4. Nolan DJ, Ginsberg M, Israely E, et al. Molecular signatures of tissue-specific microvascular endothelial cell heterogeneity in organ maintenance and regeneration. Dev Cell. 2013;26:204–219.

5. Coppiello G, Collantes M, Sirerol-Piquer MS, et al. Meox2/Tcf15 heterodimers program the heart capillary endothelium for cardiac fatty acid uptake. Circulation. 2015;131:815–826.

6. Pinto AR, Ilinykh A, Ivey MJ, et al. Revisiting Cardiac Cellular Composition. Circ Res. 2016;118:400–409.

7. Talman V, Kivelä R. Cardiomyocyte-Endothelial Cell Interactions in Cardiac Remodeling and Regeneration. Front Cardiovasc Med. 2018;5:101.

8. Olsson AK, Dimberg A, Kreuger J, Claesson-Welsh L. VEGF receptor signalling - in control of vascular function. Nat Rev Mol Cell Biol. 2006;7:359–371.

9. Mohan P, Brutsaert DL, Paulus WJ, Sys SU. Myocardial contractile response to nitric oxide and cGMP. Circulation. 1996;93:1223–1229.

10. Drawnel FM, Archer CR, Roderick HL. The role of the paracrine/autocrine mediator endothelin-1 in regulation of cardiac contractility and growth. Br J Pharmacol. 2013;168:296–317.

11. Wang W, McKinnie SMK, Patel VB, et al. Loss of Apelin exacerbates myocardial infarction adverse remodeling and ischemia-reperfusion injury: therapeutic potential of synthetic Apelin analogues. J Am Heart Assoc. 2013;2:e000249.

12. Gemberling M, Karra R, Dickson AL, Poss KD. Nrg1 is an injury-induced cardiomyocyte mitogen for the endogenous heart regeneration program in zebrafish. Elife. 2015;4.

13. Takahashi K, Yamanaka S. Induction of pluripotent stem cells from mouse embryonic and adult fibroblast cultures by defined factors. Cell. 2006;126:663–676.

14. Hamazaki T, El Rouby N, Fredette NC, Santostefano KE, Terada N. Concise Review: Induced Pluripotent Stem Cell Research in the Era of Precision Medicine. Stem Cells. 2017;35:545–550.

15. Ikuno T, Masumoto H, Yamamizu K, Yoshioka M, Minakata K, Ikeda T, Sakata R, Yamashita JK. Efficient and robust differentiation of endothelial cells from human induced pluripotent stem cells via lineage control with VEGF and cyclic AMP. PLoS One. 2017;12:e0173271.

16. Rufaihah AJ, Huang NF, Kim J, et al endothelial cells exhibit functional heterogeneity. Am J Transl Res. 2013;5:21–35.

17. Yang X, Pabon L, Murry CE. Engineering adolescence: maturation of human pluripotent stem cell–derived cardiomyocytes. Circ Res. 2014;114:511–523.

18. Trokovic R, Weltner J, Otonkoski T. Generation of hiPS C line HEL47.2 from healthy human adult fibroblasts. Stem Cell Res. 2015;15:263–265.

19. Trokovic R, Weltner J, Otonkoski T. Generation of hiPS C line HEL24.3 from human neonatal foreskin fibroblasts. Stem Cell Res. 2015;15:266–268.

20. Saarimäki-Vire J, Balboa D, Russell MA, et al. An Activating STAT3 Mutation Causes Neonatal Diabetes through Premature Induction of Pancreatic Differentiation. Cell Rep. 2017;19:281–294.

21. Hochbaum DR, Zhao Y, Farhi SL, et al. All-optical electrophysiology in mammalian neurons using engineered microbial rhodopsins. Nat Methods. 2014;11:825–833.

22. Dempsey GT, Chaudhary KW, Atwater N, Nguyen C, Brown BS, McNeish JD, Cohen AE, Kralj JM. Cardiotoxicity screening with simultaneous optogenetic pacing, voltage imaging and calcium imaging. J Pharmacol Toxicol Methods. 2016;81:240–250.

23. Björk S, Ojala EA, Nordström T, Ahola A, Liljeström M, Hyttinen J, Kankuri E, Mervaala E. Evaluation of Optogenetic Electrophysiology Tools in Human Stem Cell-Derived Cardiomyocytes. Front Physiol. 2017;8:884.

24. Butler A, Hoffman P, Smibert P, Papalexi E, Satija R. Integrating single-cell transcriptomic data across different conditions, technologies, and species. Nat Biotechnol. 2018;36:411–420.

25. Cui Y, Zheng Y, Liu X, et al. Single-Cell Transcriptome Analysis Maps the Developmental Track of the Human Heart. Cell Rep. 2019;26:1934–1950.e5.

26. Bao ZZ, Bruneau BG, Seidman JG, Seidman CE, Cepko CL. Regulation of chamber-specific gene expression in the developing heart by Irx4. Science. 1999;283:1161–1164.

27. Devalla HD, Schwach V, Ford JW, et al. Atrial-like cardiomyocytes from human pluripotent stem cells are a robust preclinical model for assessing atrial-selective pharmacology. EMBO Mol Med. 2015;7:394–410.

28. Marcu R, Choi YJ, Xue J, et al. Human Organ-Specific Endothelial Cell Heterogeneity. iScience. 2018;4:20–35.

29. Paul MH, Harvey RP, Wegner M, Sock E. Cardiac outflow tract development relies on the complex function of Sox4 and Sox11 in multiple cell types. Cell Mol Life Sci. 2014;71:2931–2945.

30. Kechele DO, Dunworth WP, Trincot CE, Wetzel-Strong SE, Li M, Ma H, Liu J, Caron KM. Endothelial Restoration of Receptor Activity-Modifying Protein 2 Is Sufficient to Rescue Lethality, but Survivors Develop Dilated Cardiomyopathy. Hypertension. 2016;68:667–677.

31. Materna SC, Sinha T, Barnes RM, Lammerts van Bueren K, Black BL. Cardiovascular development and survival require Mef2c function in the myocardial but not the endothelial lineage. Dev Biol. 2019;445:170–177.

32. Dackor RT, Fritz-Six K, Dunworth WP, Gibbons CL, Smithies O, Caron KM. Hydrops fetalis, cardiovascular defects, and embryonic lethality in mice lacking the calcitonin receptor-like receptor gene. Mol Cell Biol. 2006;26:2511–2518.

33. Deshwar AR, Chng SC, Ho L, Reversade B, Scott IC. The Apelin receptor enhances Nodal/TGFβ signaling to ensure proper cardiac development. Elife. 2016;5.

34. Wirrig EE, Snarr BS, Chintalapudi MR, et al. Cartilage link protein 1 (Crtl1), an extracellular matrix component playing an important role in heart development. Dev Biol. 2007;310:291–303.

35. Fernández-Hernando C, Yu J, Dávalos A, Prendergast J, Sessa WC. Endothelial-specific overexpression of caveolin-1 accelerates atherosclerosis in apolipoprotein E-deficient mice. Am J Pathol. 2010;177:998–1003.

36. Guo L, Zhang H, Hou Y, Wei T, Liu J. Plasmalemma vesicle-associated protein: A crucial component of vascular homeostasis. Exp Ther Med. 2016;12:1639–1644.

37. Ohashi M, Runge MS, Faraci FM, Heistad DD. MnSOD deficiency increases endothelial dysfunction in ApoE-deficient mice. Arterioscler Thromb Vasc Biol. 2006;26:2331–2336.

38. Sangwung P, Zhou G, Nayak L, et al. KLF2 and KLF4 control endothelial identity and vascular integrity. JCI Insight. 2017;2:e91700.

39. Issan Y, Kornowski R, Aravot D, Shainberg A, Laniado-Schwartzman M, Sodhi K, Abraham NG, Hochhauser E. Heme oxygenase-1 induction improves cardiac function following myocardial ischemia by reducing oxidative stress. PLoS One. 2014;9:e92246.

40. Dinkova-Kostova AT, Talalay P. Persuasive evidence that quinone reductase type 1 (DT diaphorase) protects cells against the toxicity of electrophiles and reactive forms of oxygen. Free Radic Biol Med. 2000;29:231–240.

41. Conway DE, Sakurai Y, Weiss D, Vega JD, Taylor WR, Jo H, Eskin SG, Marcus CB, McIntire LV. Expression of CYP1A1 and CYP1B1 in human endothelial cells: regulation by fluid shear stress. Cardiovasc Res. 2009;81:669–677.

42. Bhattacharya R, Wang E, Dutta SK, Vohra PK, E G, Prakash YS, Mukhopadhyay D. NHERF-2 maintains endothelial homeostasis. Blood. 2012;119:4798–4806.

43. Schrimpf C, Xin C, Campanholle G, et al. Pericyte TIMP3 and ADAMTS1 modulate vascular stability after kidney injury. J Am Soc Nephrol. 2012;23:868–883.

44. Yanagisawa M, Kurihara H, Kimura S, Tomobe Y, Kobayashi M, Mitsui Y, Yazaki Y, Goto K, Masaki T. A novel potent vasoconstrictor peptide produced by vascular endothelial cells. Nature. 1988;332:411–415.

45. Masoud AG, Lin J, Azad AK, et al. Apelin directs endothelial cell differentiation and vascular repair following immune-mediated injury. J Clin Invest. 2020;130:94–107.

46. Freedman SJ, Sun Z-YJ, Kung AL, France DS, Wagner G, Eck MJ. Structural basis for negative regulation of hypoxia-inducible factor-1α by CITED2. Nat Struct Mol Biol. 2003;10:504–512.

47. Trittmann JK, Almazroue H, Jin Y, Nelin LD. DDAH1 regulates apoptosis and angiogenesis in human fetal pulmonary microvascular endothelial cells. Physiol Rep. 2019;7:e14150.

48. Smadja DM, d’Audigier C, Bièche I, et al. Thrombospondin-1 is a plasmatic marker of peripheral arterial disease that modulates endothelial progenitor cell angiogenic properties. Arterioscler Thromb Vasc Biol. 2011;31:551–559.

49. Adams GN, Stavrou EX, Fang C, et al. Prolylcarboxypeptidase promotes angiogenesis and vascular repair. Blood. 2013;122:1522–1531.

50. Kinderlerer AR, Ali F, Johns M, et al. KLF2-dependent, shear stress-induced expression of CD59: a novel cytoprotective mechanism against complement-mediated injury in the vasculature. J Biol Chem. 2008;283:14636–14644.

51. Kern CB, Wessels A, McGarity J, et al. Reduced versican cleavage due to Adamts9 haploinsufficiency is associated with cardiac and aortic anomalies. Matrix Biol. 2010;29:304–316.

52. Torry RJ, Tomanek RJ, Zheng W, Miller SJ, Labarrere CA, Torry DS. Hypoxia increases placenta growth factor expression in human myocardium and cultured neonatal rat cardiomyocytes. J Heart Lung Transplant. 2009;28:183–190.

53. Accornero F, van Berlo JH, Benard MJ, Lorenz JN, Carmeliet P, Molkentin JD. Placental growth factor regulates cardiac adaptation and hypertrophy through a paracrine mechanism. Circ Res. 2011;109:272–280.

54. Paik DT, Tian L, Lee J, et al. Large-Scale Single-Cell RNA-Seq Reveals Molecular Signatures of Heterogeneous Populations of Human Induced Pluripotent Stem Cell-Derived Endothelial Cells. Circ Res. 2018;123:443–450.

55. Vilà-González M, Kelaini S, Magee C, et al. Enhanced Function of Induced Pluripotent Stem Cell-Derived Endothelial Cells Through ESM1 Signaling. Stem Cells. 2019;37:226–239.

56. Ohtani-Kaneko R, Sato K, Tsutiya A, Nakagawa Y, Hashizume K, Tazawa H. Characterisation of human induced pluripotent stem cell-derived endothelial cells under shear stress using an easy-to-use microfluidic cell culture system. Biomed Microdevices. 2017;19:91.

57. Wu S-P, Cheng C-M, Lanz RB, et al. Atrial identity is determined by a COUP-TFII regulatory network. Dev Cell. 2013;25:417–426.

58. Barnes RM, Firulli BA, Conway SJ, Vincentz JW, Firulli AB. Analysis of the Hand1 cell lineage reveals novel contributions to cardiovascular, neural crest, extra-embryonic, and lateral mesoderm derivatives. Dev Dyn. 2010;239:3086–3097.

59. de Pater E, Ciampricotti M, Priller F, et al. Bmp signaling exerts opposite effects on cardiac differentiation. Circ Res. 2012;110:578–587.

60. MacGrogan D, D’Amato G, Travisano S, et al. Sequential Ligand-Dependent Notch Signaling Activation Regulates Valve Primordium Formation and Morphogenesis. Circ Res. 2016;118:1480–1497.

61. D’Amato G, Luxán G, del Monte-Nieto G, et al. Sequential Notch activation regulates ventricular chamber development. Nat Cell Biol. 2016;18:7–20.

62. Luxán G, D’Amato G, MacGrogan D, de la Pompa JL. Endocardial Notch Signaling in Cardiac Development and Disease. Circ Res. 2016;118:e1–e18.

63. Loomes KM, Underkoffler LA, Morabito J, Gottlieb S, Piccoli DA, Spinner NB, Baldwin HS, Oakey RJ. The expression of Jagged1 in the developing mammalian heart correlates with cardiovascular disease in Alagille syndrome. Hum Mol Genet. 1999;8:2443–2449.

64. Li Z, Solomonidis EG, Meloni M, et al. Single-cell transcriptome analyses reveal novel targets modulating cardiac neovascularization by resident endothelial cells following myocardial infarction. Eur Heart J. 2019;40:2507–2520.

65. Williams IM, Wu JC. Generation of Endothelial Cells From Human Pluripotent Stem Cells. Arterioscler Thromb Vasc Biol. 2019;39:1317–1329.

66. Giacomelli E, Bellin M, Orlova VV, Mummery CL. Co-Differentiation of Human Pluripotent Stem Cells-Derived Cardiomyocytes and Endothelial Cells from Cardiac Mesoderm Provides a Three-Dimensional Model of Cardiac Microtissue. Curr Protoc Hum Genet. 2017;95:21.9.1-21.9.22.

67. Sharma A, Li G, Rajarajan K, Hamaguchi R, Burridge PW, Wu SM. Derivation of highly purified cardiomyocytes from human induced pluripotent stem cells using small molecule-modulated differentiation and subsequent glucose starvation. J Vis Exp. 2015.

68. Butler, A., Hoffman, P., Smibert, P., Papalexi, E. & Satija, R. Integrating single-cell transcriptomic data across different conditions, technologies, and species. Nat. Biotechnol. 2018; 36, 411–420.

69. Hafemeister C, Satija R. Normalization and variance stabilization of single-cell RNA-seq data using regularized negative binomial regression. Genome Biol. 2019;20:296.

