## Supplemental Data for "Modeling cellular crosstalk and organotypic vasculature development with human iPSC-derived endothelial cells and cardiomyocytes"

Supplemental material

### Supplemental methods

*iPSC Culture*
iPSCs were maintained in Essential 8 media (A1517001, Thermo Fisher Scientific) on thin-coated Matrigel (354277, dilution 1:200; Corning, Corning, NY). The cells were passaged using EDTA.

*iPS-EC differentiation*
Endothelial cell differentiation was conducted based on the protocol by Giacomelli et al.[66]. The BPEL medium ingredients were purchased from the same vendors as mentioned in the article, except for BSA (A7030, Sigma) and PVA (362607, Sigma). Briefly, 125 000 - 175 000 cells/well in a 6-well plate were plated on day 0. On day 1, the medium was changed to BPEL with 20 ng/ml BMP4 (120-05ET, Peprotech), 20 ng/ml Activin A (AF-120-14E-50ug, Peprotech) and 4 µmol/L CHIR (S2924, Selleckhem). On day 3, the medium was changed to BPEL with 50 ng/ml VEGF (produced in-house) and 5 µmol/L IWR-1 (I0161, Sigma). On day 6, medium was changed to BPEL with 50 ng/ml VEGF and the cells were maintained in this medium until they were sorted. 50 ng/ml VEGF was maintained in all cell culture with iPS-ECs unless otherwise indicated.

*iPSC-EC sorting*
After differentiation, iPS-ECs were sorted using magnetic beads with an antibody against CD31 (130-091-935, Miltenyi Biotec), according to the manufacturer's protocol. The concentration of the cells was counted with Bio-Rad TC10 or TC20 Automated Cell Counter. The cells were immediately used for experiments.

*IPS-CM differentiation*
iPS-CM differentiation was modified from Sharma et al.[67]. iPS cells were plated 1:10 -1:15 in 12-well plates coated with Matrigel. After reaching confluency (day 0), differentiation was started by changing the media to RPMI (10-040-CV, Corning) and B27 supplement without insulin (A1895601, Thermo Fisher Scientific), with 5 µmol/L CHIR. The following day (day 1), the same media with CHIR was added to the cells. On day 2, the media was changed for RPMI/B27 without insulin, with 5 µmol/L IWR-1, and replenished on day 3. On day 4 and 5, RPMI/B27 without insulin was used. For days 6 to 8, the media was changed to RPMI media and B27 with insulin (17504044, Thermo Fisher Scientific). On day 9, RPMI without glucose (10-043-CV, Corning) and B27 with insulin, but with added lactate (L7900, Sigma, 466 µl/500ml) was used. The cells were maintained in this media until they were used in experiments. The media was changed to BPEL two to three days before the start of the experiments.

*qPCR analysis of CM marker gene expression*
RNA samples were collected from CM differentiation experiments on days 1, 3, 5, 7, 9, 11, 13 and 20. RNA was extracted using Nucleospin RNA Plus Extraction kit (740984, Macherey-Nagel). cDNA synthesis was performed using High Capacity cDNA Reverse Transcription Kit (4368814, Thermo) qPCR analysis was done using SYBR Green (04913914001, Roche) and Bio-Rad CFX96 Real-Time PCR Detection System. All results were first normalized to housekeeping gene RPL37A and then to the expression levels during day 1.

*Immunofluorescence staining*
Cells were fixed with 4% PFA and stained with monoclonal anti-α-Actinin (Sarcomeric) (A7811, Sigma) and VE-cadherin (2500, Cell Signaling Technology) antibodies. Nuclei were visualized with

DAPI or Hoechst. Stained cells were imaged with fluorescent or confocal microscopes (Zeiss AxioImager and Zeiss LSM 780).

*Matrigel tube assay*
48-well plate was coated with 100 µl of Matrigel per well. After gelling of Matrigel, 90 000 iPS-ECs/well (HEL24.3, HEL47.2) or 30 000 HUVECs/well were added on top of the Matrigel-covered wells. The cells were allowed to attach and grow. Phase-contrast images were taken at 24, 48 and 72 hours.

*LDL uptake*
Atto LDL oxidized with 10 µmol/L $CuSO_4$ (20h, 37C°) (courtesy of Katariina Öörni lab) was applied to ECs on coverslips in a 24-well plate for 20h (7.5 µg/well). The coverslips were fixed and imaged with a Zeiss LSM 780 confocal microscope.

*Preparation of iPS-CMs for optogenetics*
For optogenetic analysis, lentiviral vector Optopatch (a kind gift from Adam E. Cohen group,[21,22] acquired through Addgene, plasmid # 62984), was introduced in iPS-CMs, using 1 pg of virus per cell for transduction. During lentiviral work, the cells were cultured in 12-well plates coated with Matrigel and detached twice using Accutase to remove replicating virus. RPMI/B27 medium with glucose was used 48 hours after transduction and both passages. Otherwise the cells were cultured in RPMI/B27 medium with lactate. On the second passage, the cells were plated on Matrigel-coated glass-bottom dishes (14 mm Ø, P35G-1.5-14-C, MatTek) for optogenetic imaging. William's E medium (Life Technologies A12176) supplemented with Cocktail B (Life Technologies CM4000) without dexamethasone was used as imaging medium.

*Optogenetic video imaging of iPS-CMs*
Action potentials were recorded from spontaneously beating iPS-CMs expressing Optopatch. Optogenetic imaging platform included environmental chamber (5% $CO_2$, 37°C, EMBL), Nikon Eclipse Ti-E fluorescence microscope, red laser light source ($\lambda$ = 647 nm, Ef = 550 mW/mm2 ) and NIS-Elements advanced research software for the platform operation. Raw data was recorded as image sequences (50 frames per second), from which the total fluorescence intensity signal was exported to MS Excel. Data was normalized by fitting the acquired signal to an exponential function with cPot Cardiac Action Potential Calculator software, written in MATLAB. The optogenetic imaging platform and workflow is described in more detail in Björk et al., 2017[23].

*Single-cell sequencing*
Single-cell gene expression profiles were studied using the 10x Genomics Chromium Single Cell 3'RNAseq platform. The Chromium Single Cell 3'RNAseq run and library preparation were done using the Chromium™ Single Cell 3' Reagent version 2 chemistry. The sample libraries were sequenced on Illumina NovaSeq 6000 system. 4000 cells and 50 000 PE/cell were analyzed.

Data processing and analysis were performed using 10x Genomics Cell Ranger v2.1.1 pipelines. Cell Ranger includes several pipelines of which the "cellranger mkfastq" was used to produce FASTQ (raw data) files and "cellranger count" to perform alignment, filtering and UMI counting. mkfastq was run using the Illumina bcl2fastq v2.2.0 and alignment was done against human genome GRCh38. Cellranger aggr pipeline was used to combine data from multiple samples into an experiment-wide gene-barcode matrix and analysis.

Analyses were performed with Seurat R package version 3.0.1[68]. Cells, in which more than 1000 genes were detected, were included. Seurat function CellCycleScoring was used to assign cell cycle scores (iG2/M scores and S scores), which then were used to regress out cell cycle effect. Normalization and variance stabilization was done with SCTransform in Seurat (vars.to.regress was used to remove confounding sources of variation including mitochondrial mapping percentage, and cell cycle scores)[69]. Principal component analysis (PCA) was performed on the highly variable genes, and the first 30 PCs were used for uniform manifold approximation (UMAP). Differential expression for each subpopulation in the scRNA-seq data was performed using the Wilcoxon Rank Sum test by FindAllMarkers function in Seurat, and FindMarkers was used to distinguish different conditions. Cells were clustered based on their expression profile. Cell types were annotated based on established markers on The Human Protein Atlas and published literature.

*RNA-sequencing*

RNA samples were sequenced with Illumina NextSeq sequencer (Illumina, San Diego, CA, USA) in High output run using NEBNext® Ultra™ II Directional RNA Library Prep Kit for Illumina. The sequencing was performed as single-end sequencing for read length 75 bp. The count data was used to calculate differential expression statistics with the DESeq2 software in R environment. Genes with an adjusted p-value for the log2-fold change $< 0.05$ were considered significant.

Figure I: Expression of cardiomyocyte markers during differentiation. qPCR analysis shows that CM marker gene expression is increased during differentiation. Two differentiation experiments (HEL47.2 (1.) and HEL47.2 (2.)) with two replicates each are shown. Expression levels have been normalized by comparing to day 1

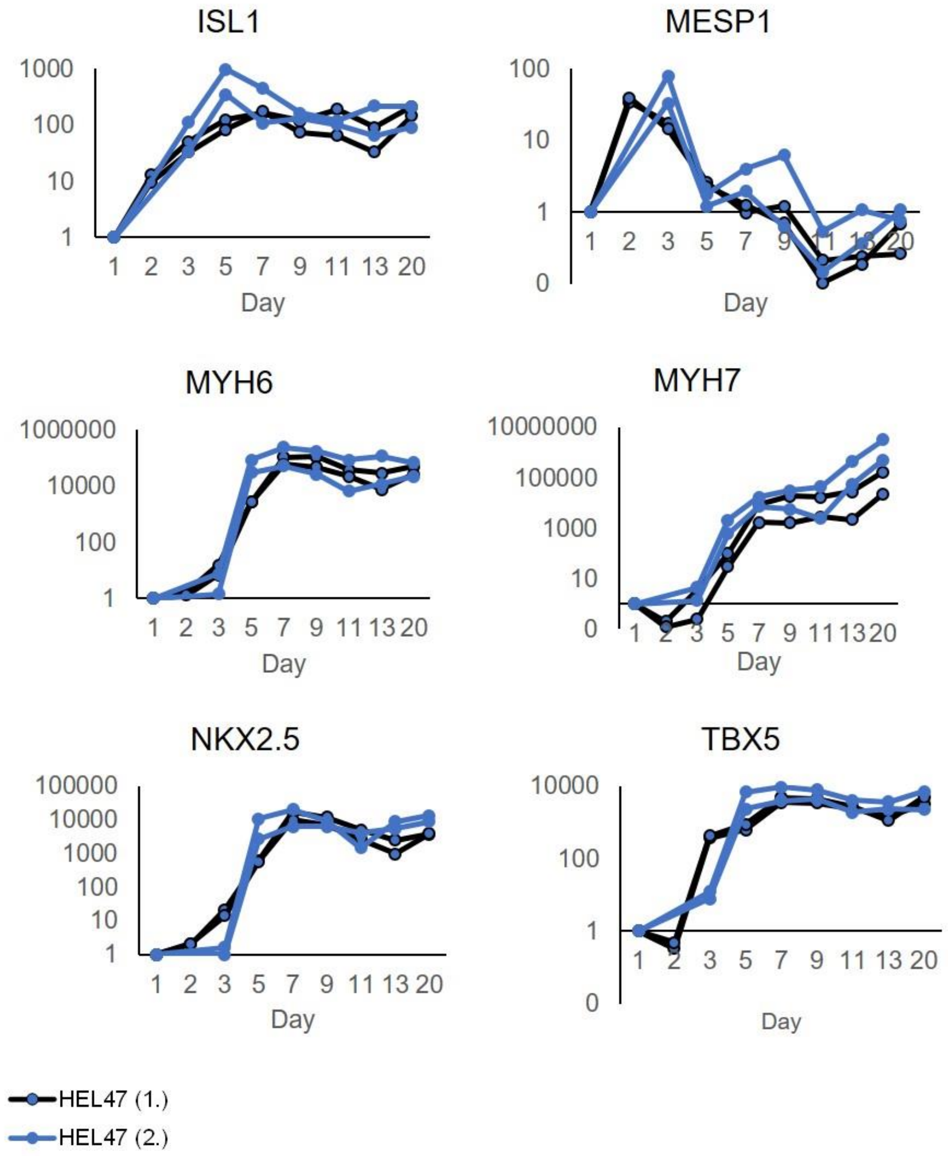

Figure II: **: A,** Sample wise UMAP clustering of HEL47.2. **B,** Distributions of sample cells per cluster.

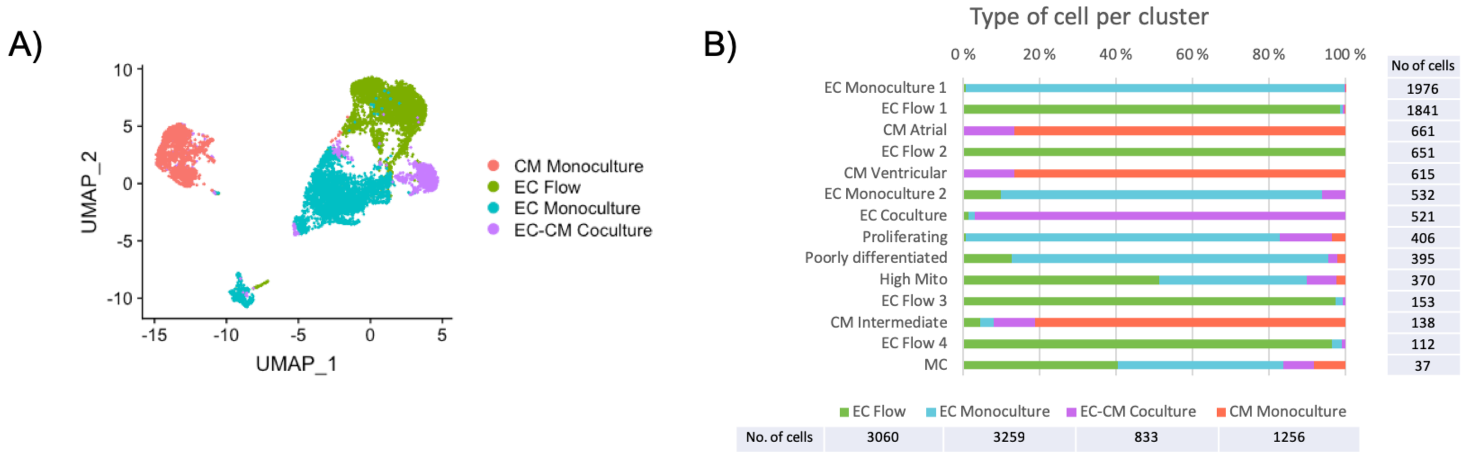

Figure III: Clustering of HEL24.3 cells and HEL47.2 without flow cells.

HEL 24.3

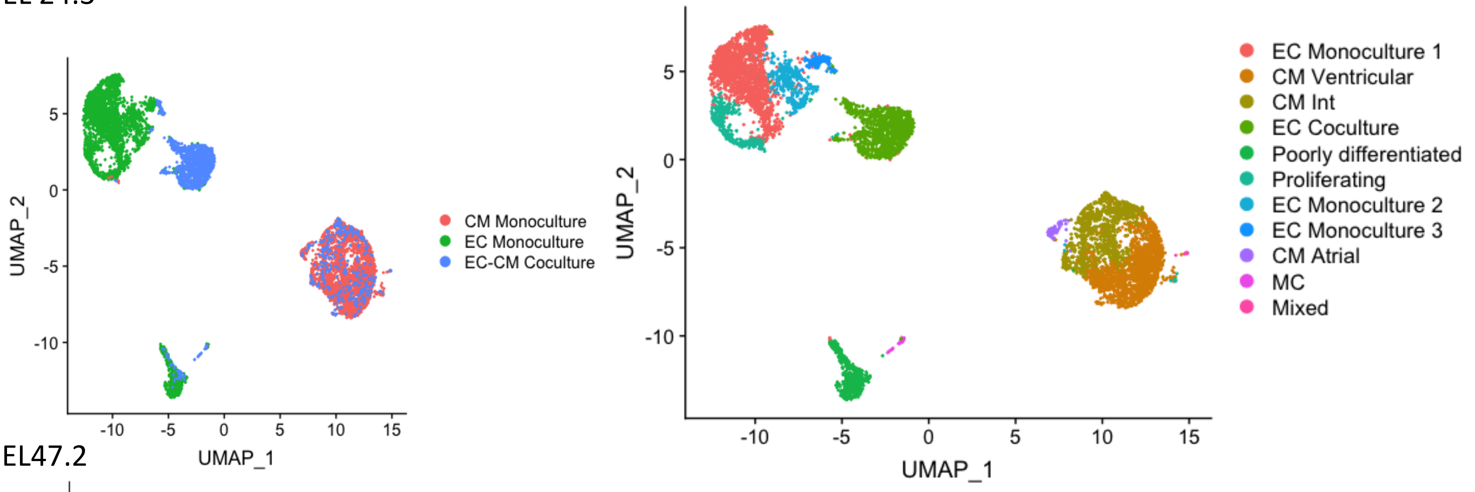

HEL47.2

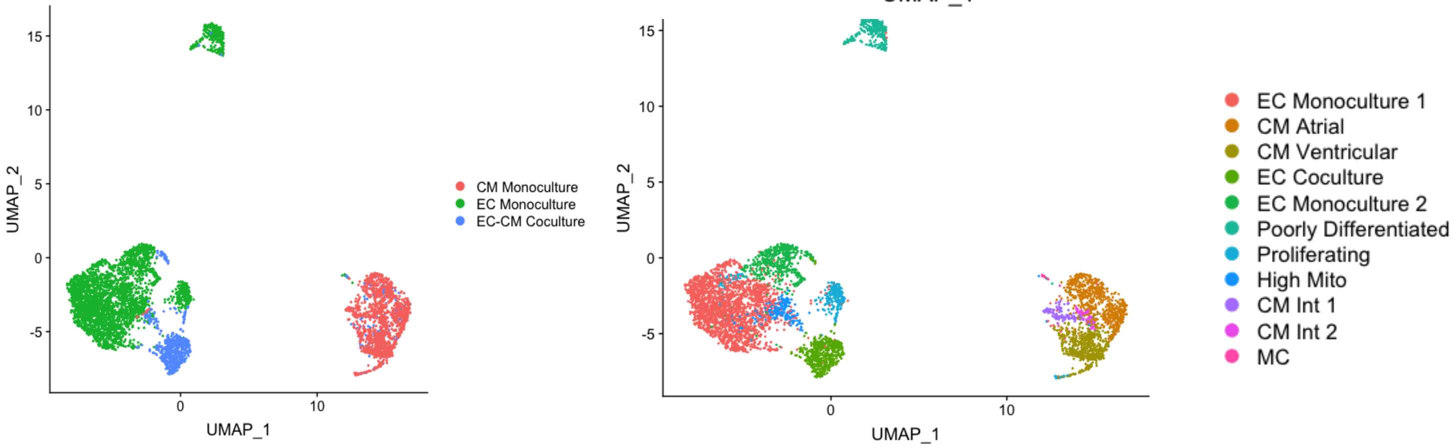

Figure IV: The expression of A, endothelial markers, B, cardiomyocyte markers in HEL24.3 iPS ECs. The expression of C, ventricular and, D, atrial genes in HEL24.3 iPS-CMs.

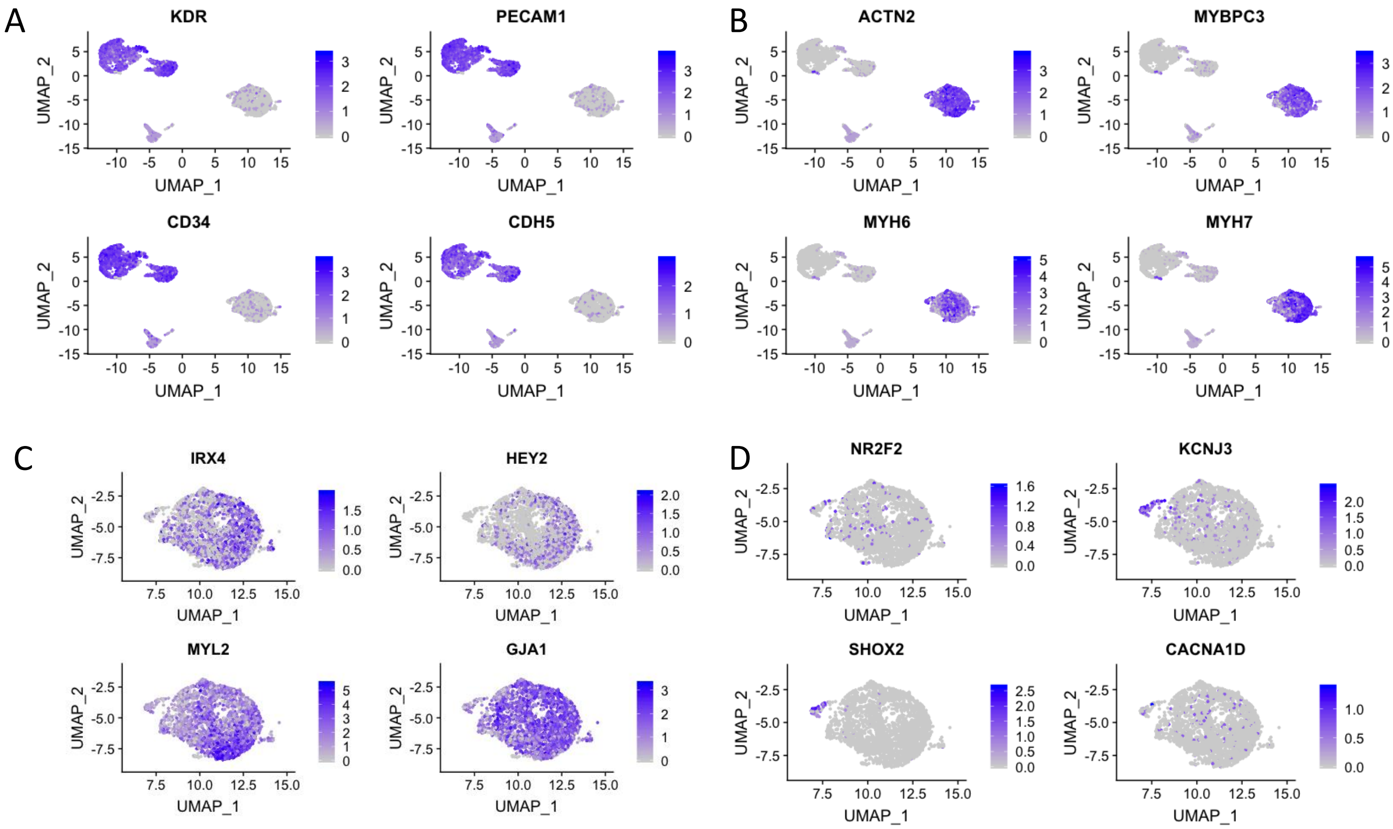

Figure V: The expression of cardiac specific EC-genes, genes associated with heart development, and genes associated with regulation of angiogenesis in HEL24.3 monoculture and coculture iPS-ECs (all differences between EC Coculture and EC Monoculture are statistically significant, adjusted P<0.05).

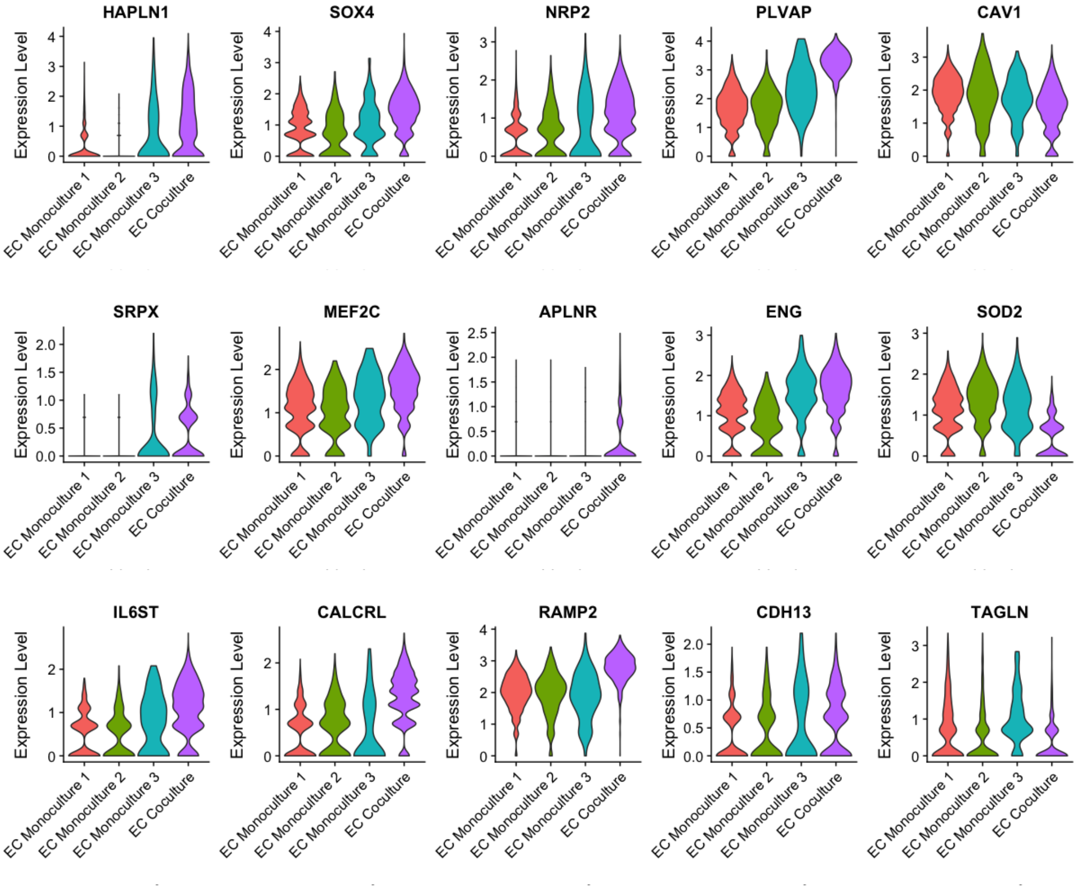

Figure VI: The expression of BMP pathway genes in HEL24.3 monoculture and coculture iPS-ECs and iPS-CMs. * = Significantly higher expression in coculture vs. monoculture cells (adjusted p<0.05), **=Significantly lower expression in coculture vs. monoculture cells (adjusted p<0.05).

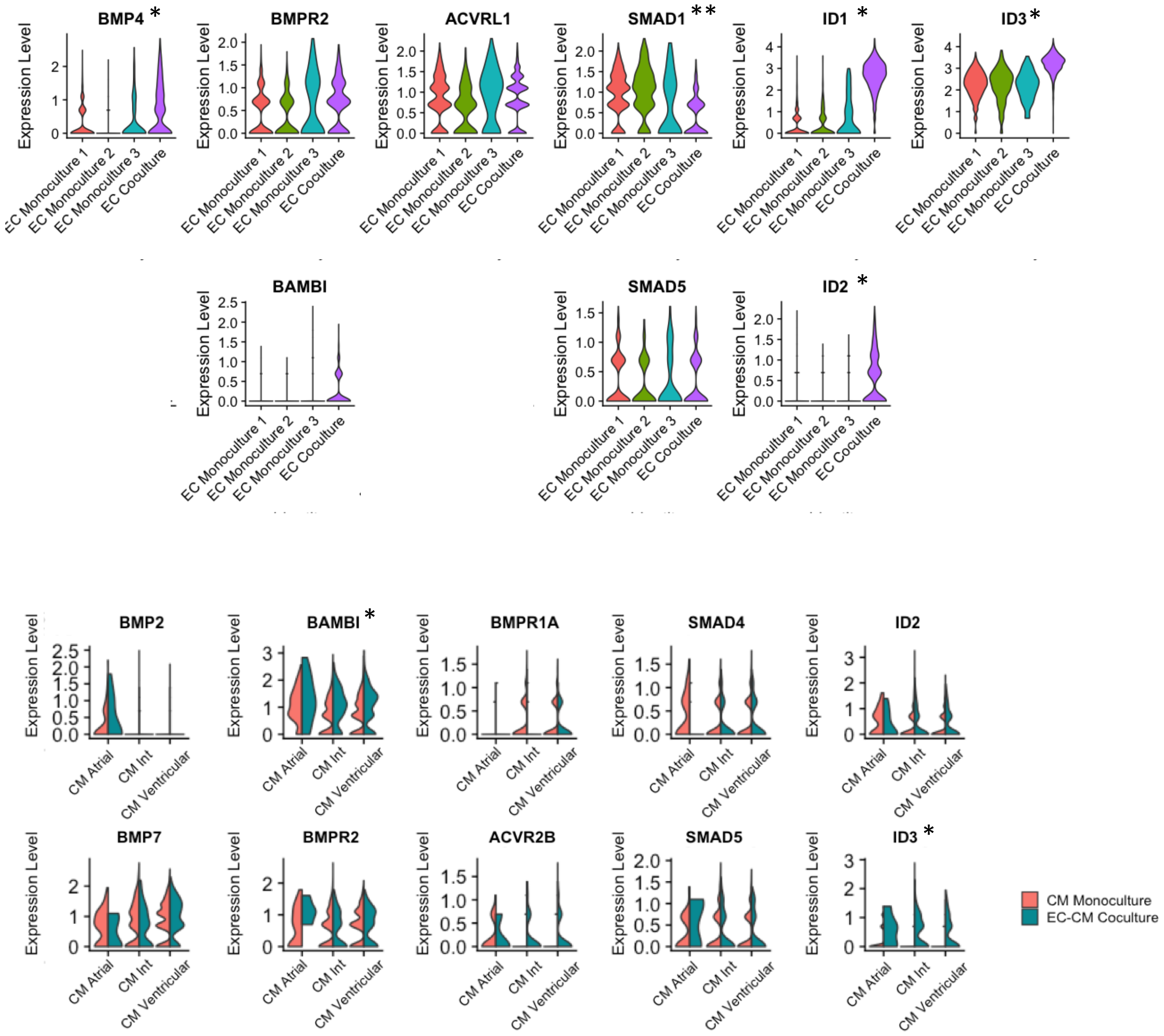

Figure VII: The expression of NOTCH-pathway genes in HEL47.2 and HEL24.3 monoculture and coculture iPS-ECs and iPS-CMs. * = Significantly higher expression in coculture vs. monoculture cells (adjusted p<0.05),

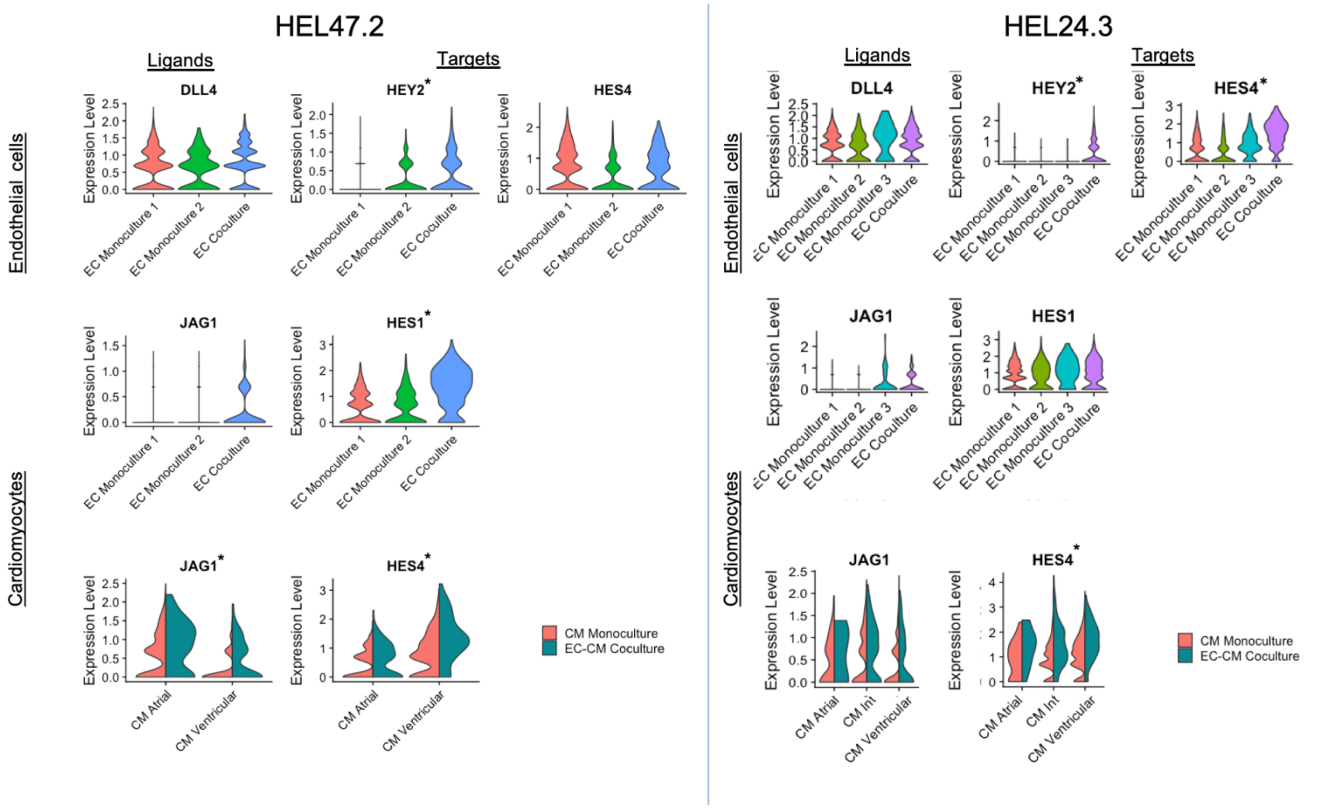

Figure VIII: **A,** Clustering of HEL24.3 static EC Monoculture and EC Flow cells. The expression of **B,** venous and **C,** arterial genes in these cells. EC Flow cluster 5 has an arterial-like gene expression profile.

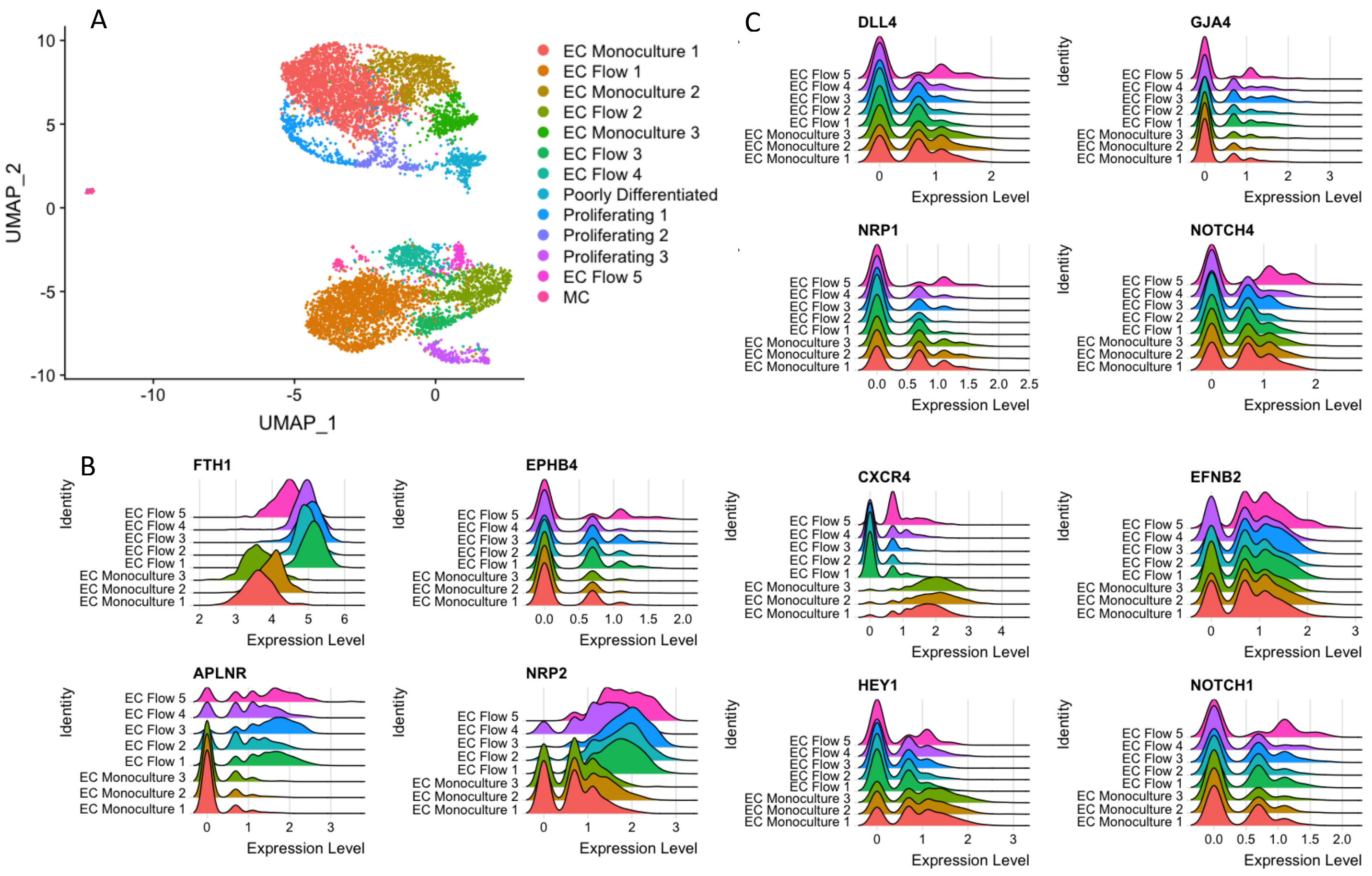

Figure IX: Top 25 up- and downregulated genes in flow in the four iPS-EC cell lines analysed by RNASeq.

| HEL46.11 Flow | HEL47.2 Flow | HEL24.3 Flow | K1 Flow | HEL46.11 Static | HEL47.2 Static | HEL24.3 Static | K1 Static | Gene |
|---|---|---|---|---|---|---|---|---|
| 12.322 | 11.616 | 12.039 | 12.246 | 8.494 | 7.930 | 8.192 | 8.457 | KLF2 (ENSG00000127528.5) |
| 11.143 | 9.978 | 10.181 | 10.435 | 7.160 | 6.469 | 6.512 | 6.889 | KLF4 (ENSG00000136826.15) |
| 12.595 | 12.365 | 12.287 | 12.811 | 10.918 | 11.045 | 10.938 | 11.033 | PTPRG (ENSG00000144724.20) |
| 8.848 | 8.292 | 8.534 | 9.318 | 5.801 | 5.953 | 5.605 | 5.803 | OPN5 (ENSG00000124818.15) |
| 13.414 | 13.274 | 13.014 | 13.599 | 10.298 | 10.678 | 10.697 | 11.326 | SLC9A3R2 (ENSG00000065054.14) |
| 10.963 | 10.418 | 10.716 | 10.902 | 8.935 | 8.490 | 8.697 | 8.724 | TSC22D3 (ENSG00000157514.16) |
| 10.902 | 9.543 | 10.829 | 11.290 | 7.287 | 6.975 | 7.885 | 7.073 | HSPA12B (ENSG00000132622.11) |
| 11.028 | 10.401 | 10.951 | 10.853 | 8.583 | 8.792 | 8.736 | 9.450 | TLR4 (ENSG00000136869.15) |
| 15.438 | 15.294 | 15.100 | 15.274 | 12.294 | 13.194 | 12.992 | 13.164 | PODXL (ENSG00000128567.17) |
| 13.574 | 13.585 | 13.318 | 13.434 | 11.109 | 12.093 | 11.441 | 11.422 | HEG1 (ENSG00000173706.14) |
| 10.389 | 10.454 | 10.426 | 10.436 | 9.909 | 9.917 | 9.969 | 9.994 | FBXO30 (ENSG00000118496.5) |
| 11.433 | 10.926 | 11.702 | 11.247 | 9.414 | 9.844 | 9.383 | 9.508 | DYRK2 (ENSG00000127334.10) |
| 12.865 | 12.465 | 12.643 | 12.752 | 11.316 | 11.436 | 10.805 | 11.080 | HYAL2 (ENSG00000068001.14) |
| 13.412 | 12.971 | 13.142 | 13.114 | 12.190 | 12.182 | 12.102 | 11.933 | HIPK2 (ENSG00000064393.16) |
| 9.167 | 8.822 | 8.084 | 9.776 | 6.306 | 6.134 | 5.822 | 6.455 | CMKLR1 (ENSG00000174600.14) |
| 10.537 | 9.890 | 10.144 | 10.179 | 9.199 | 9.089 | 8.966 | 8.973 | CARD10 (ENSG00000100065.15) |
| 12.325 | 12.316 | 12.066 | 12.426 | 11.508 | 11.467 | 11.561 | 11.636 | SPAG9 (ENSG00000008294.21) |
| 8.840 | 8.360 | 9.328 | 9.572 | 6.550 | 6.375 | 7.035 | 6.848 | CFAP161 (ENSG00000156206.14) |
| 10.056 | 9.805 | 9.453 | 10.009 | 8.067 | 8.360 | 8.423 | 8.517 | MGAT4A (ENSG00000071073.13) |
| 10.095 | 9.827 | 9.765 | 9.896 | 8.776 | 9.048 | 8.963 | 8.851 | KLF10 (ENSG00000155090.15) |
| 10.749 | 10.541 | 10.201 | 10.917 | 9.348 | 9.053 | 8.621 | 8.868 | ENDOD1 (ENSG00000149218.5) |
| 9.156 | 7.348 | 7.018 | 9.123 | 5.605 | 5.806 | 5.822 | 5.884 | PI16 (ENSG00000164530.15) |
| 9.743 | 9.525 | 9.506 | 9.739 | 8.604 | 8.734 | 8.666 | 8.908 | KLF11 (ENSG00000172059.11) |
| 9.434 | 8.934 | 9.340 | 9.460 | 8.130 | 8.390 | 8.044 | 8.328 | SLC8B1 (ENSG00000089060.11) |
| 12.424 | 13.739 | 10.449 | 14.417 | 6.084 | 7.538 | 6.134 | 6.698 | IGFBP5 (ENSG00000115461.5) |
| 10.857 | 10.955 | 11.168 | 10.862 | 11.996 | 12.109 | 12.216 | 11.970 | RALA (ENSG00000006451.8) |
| 7.919 | 7.804 | 7.808 | 7.900 | 8.690 | 8.475 | 8.524 | 8.357 | DALRD3 (ENSG00000178149.17) |
| 7.867 | 7.757 | 8.406 | 7.444 | 10.182 | 10.975 | 9.435 | 10.467 | BMP4 (ENSG00000125378.16) |
| 6.499 | 6.781 | 6.529 | 6.536 | 8.026 | 7.870 | 8.404 | 8.574 | PDE7B (ENSG00000171408.14) |
| 9.666 | 9.638 | 9.904 | 9.417 | 10.883 | 10.525 | 10.481 | 10.625 | NME4 (ENSG00000103202.13) |
| 8.454 | 9.799 | 10.853 | 8.407 | 12.132 | 13.372 | 12.987 | 13.066 | EDN1 (ENSG00000078401.7) |
| 8.743 | 8.704 | 9.635 | 8.774 | 10.464 | 10.840 | 10.744 | 10.790 | PDE4B (ENSG00000184588.18) |
| 8.477 | 7.750 | 8.522 | 8.175 | 10.900 | 10.149 | 9.813 | 10.039 | MAP2K6 (ENSG00000108984.15) |
| 9.144 | 9.239 | 9.192 | 9.130 | 9.684 | 9.708 | 9.640 | 9.888 | DYNLT1 (ENSG00000146425.11) |
| 7.466 | 7.777 | 7.765 | 7.805 | 8.245 | 8.868 | 8.646 | 9.022 | PIK3CD (ENSG00000171608.15) |
| 8.739 | 8.662 | 8.795 | 8.578 | 9.416 | 9.116 | 9.274 | 9.256 | SLC11A2 (ENSG00000110911.15) |
| 9.447 | 9.299 | 9.438 | 9.424 | 9.852 | 9.840 | 9.740 | 9.996 | DCTD (ENSG00000129187.14) |
| 8.936 | 9.114 | 9.300 | 9.125 | 9.934 | 9.872 | 9.801 | 9.753 | TADA3 (ENSG00000171148.13) |
| 9.160 | 9.306 | 9.370 | 9.106 | 10.040 | 10.098 | 10.115 | 10.005 | GMFB (ENSG00000197045.13) |
| 9.139 | 9.142 | 9.548 | 9.365 | 10.055 | 10.239 | 10.099 | 10.114 | RNF41 (ENSG00000181852.17) |
| 7.292 | 7.804 | 8.229 | 8.093 | 9.439 | 9.139 | 9.832 | 9.402 | PSTPIP2 (ENSG00000152229.18) |
| 10.984 | 10.818 | 11.385 | 10.763 | 12.212 | 12.095 | 12.293 | 11.830 | FAM43A (ENSG00000185112.5) |
| 6.435 | 6.103 | 7.191 | 6.454 | 9.076 | 7.918 | 9.044 | 8.619 | CALHM4 (ENSG00000164451.13) |
| 10.513 | 10.315 | 10.765 | 10.377 | 11.130 | 11.238 | 11.519 | 11.561 | RANGAP1 (ENSG00000100401.19) |
| 7.522 | 7.978 | 7.862 | 7.661 | 8.858 | 9.139 | 8.385 | 8.702 | IKBKE (ENSG00000263528.8) |
| 8.485 | 8.721 | 8.424 | 8.488 | 9.210 | 9.522 | 9.300 | 9.182 | CYB561 (ENSG00000008283.16) |
| 7.174 | 7.319 | 7.247 | 7.209 | 8.106 | 8.009 | 7.899 | 8.743 | C20orf27 (ENSG00000101220.17) |
| 11.042 | 11.310 | 11.843 | 10.481 | 12.390 | 13.034 | 12.551 | 12.745 | PDGFB (ENSG00000100311.16) |
| 9.641 | 10.338 | 10.261 | 10.283 | 10.948 | 11.633 | 11.413 | 11.535 | SLITRK4 (ENSG00000179542.16) |
| 11.977 | 12.507 | 12.567 | 12.059 | [illegible] | [illegible] | 13.137 | 13.592 | CRIM1 (ENSG00000150938.10) |

Figure X: The expression of known flow-induced genes, antiatherogenic genes, genes that promote vascular homeostasis, vascular tone regulators, and angiogenesis genes in the HEL24.3 static monoculture and flow iPS-ECs. (All differences between EC Flow and EC Monoculture are statistically significant, adjusted P<0.05)

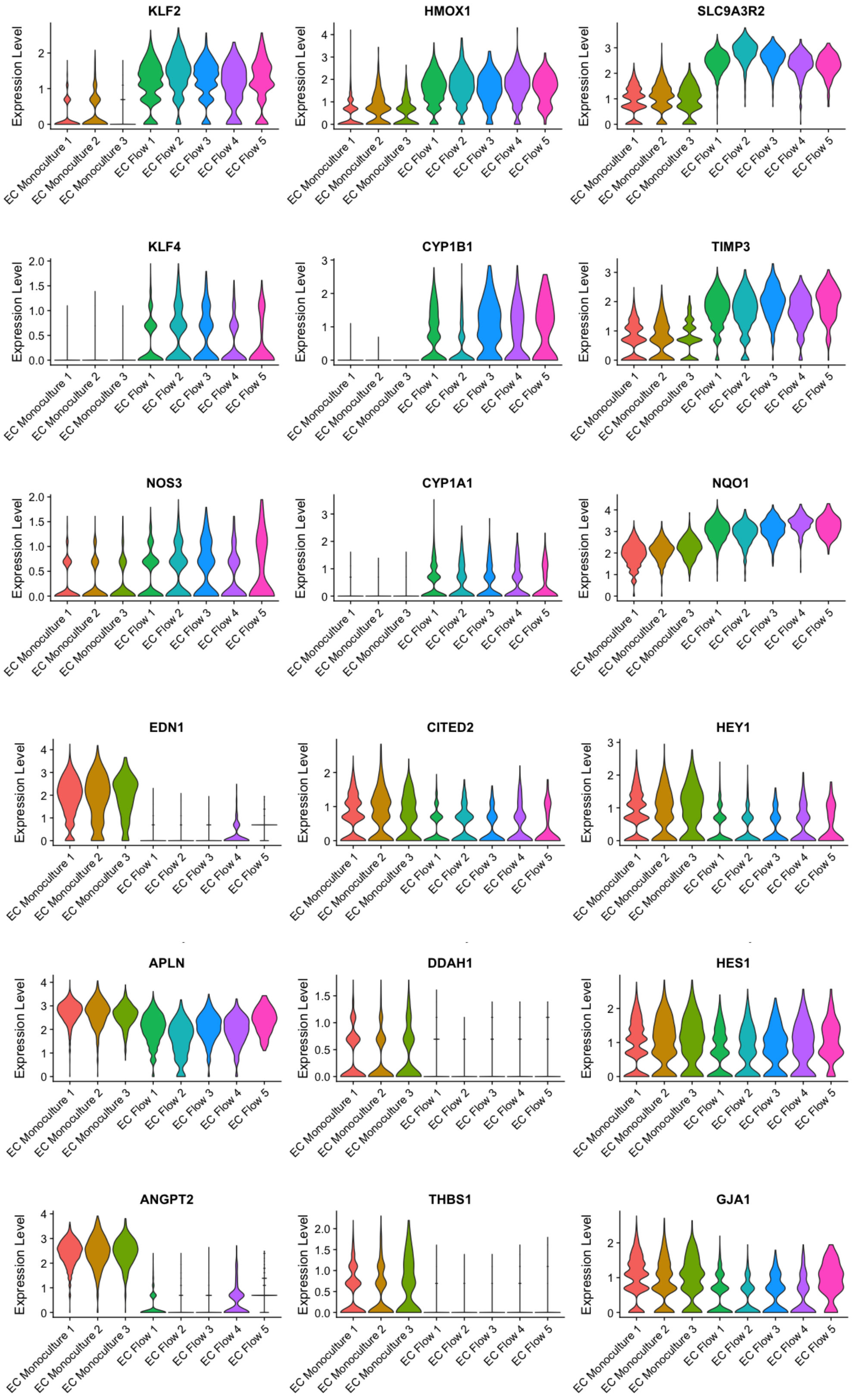

Table I: qPCR primers

| GENE | Forward primer | Reverse primer |
|---|---|---|
| ISL1 | 5'-AAAGTTACCAGCCACCTTGGA-3' | 5'-ATTAGAGCCCGGTCCTCCTT-3' |
| MESP1 | 5'-TAGGGCTAGACACTTTGAG-3' | 5'-TTTTTGCCAACTGACACC-3' |
| MYH6 | 5'-CCAGGTCAACAAGCTTCGAG-3' | 5'-TGTCACTCCTCATCGTGCAT-3' |
| MYH7 | 5'-AGTCCCAGGTCAACAAGCTG-3' | 5'-GGGCTGAGCAGATCAAGATG-3' |
| NKX2.5 | 5'-CAAGTGTGCGTCTGCCTTT-3' | 5'-TTGTCCGCCTCTGTCTTCTC-3' |
| RPL37A | 5'-GTGGTTCCTGCATGAAGACAGTG-3' | 5'-TTCTGATGGCGGACTTTACCG-3' |
| TBX5 | 5'-GGGCAGTGATGACATGGAG-3' | 5'-GCTGCTGAAAGGACTGTGGT-3' |

Table II: Upregulated and downregulated genes in iPS-ECs in coculture.

| UPREGULATED IN COCULTURE IPS-EC | | | DOWNREGULATED IN COCULTURE IPS-EC | | |
|---|---|---|---|---|---|
| **HEL47** | **HEL24_3** | **BOTH** | **HEL47** | **HEL24_3** | **BOTH** |
| ID1 | ID1 | FSTL1 | PGF | PGF | PRND |
| MT-ND3 | CYBA | PRCP | C10orf10 | C10orf10 | CREB3L1 |
| RPS27 | PLVAP | NPPA | FAM89A | FAM89A | TAGLN |
| PLVAP | HPGD | ENG | CHST1 | PHLDA1 | RPS2 |
| RPS29 | SPP1 | VAMP8 | GPX3 | TP53I11 | CXCR4 |
| PRCP | MYL7 | IER2 | TAGLN | LINC01480 | ABCA9-AS1 |
| RPL38 | GSTO1 | CD151 | SERPINB4 | CHST1 | CBLN1 |
| CYBA | VAMP5 | CYBA | PHLDA1 | CD24 | RPLP0 |
| RPL37A | PRCP | MAP1LC3A | KRT18 | SLC12A2 | EFCAB14 |
| RPL37 | HAPLN1 | RDX | RPS2 | SERPINB4 | SOD2 |
| ATP5E | CA2 | LPAR6 | EFCAB14 | FN1 | HOMER3 |
| NDUFA3 | ACTC1 | CTSB | TP53I11 | SOD2 | RP11-322E11.5 |
| NDUFB1 | TMEM88 | COL6A2 | FASN | CBLN1 | NETO2 |
| ATP5I | FABP4 | ID3 | RPS3 | CREB3L1 | LXN |
| ANKRD1 | MDK | FLT1 | MYL9 | SLC38A1 | FADS1 |
| HAPLN1 | ITM2B | PLPP3 | MGST1 | S100A6 | MT-CO2 |
| ROMO1 | ID3 | ADGRL4 | SEPW1 | ABCA9-AS1 | FASN |
| RPS28 | FCGRT | TMEM204 | TPM2 | SH3BP5 | GFOD1 |
| FCN3 | A2M | COTL1 | RPLP0 | SCD | SCD |
| SLIRP | ESM1 | WWTR1 | SSR3 | SSR3 | PGF |
| SOX4 | TNNC1 | GSN | SOD2 | TXNRD2 | MT-ATP6 |
| USMG5 | HES4 | RPS27L | RPSA | ACKR3 | RPS10 |
| HPGD | LBH | FCGRT | MVD | TUBA1B | SRM |
| TMEM258 | FLT1 | IFITM3 | NETO2 | OGFRL1 | PPP1R14B |
| RPL39 | RAMP2 | DAB2 | FAM213A | NES | SEPW1 |
| UBL5 | FCN3 | FXYD6 | NUDT4 | SMAD1 | EIF2B2 |
| RPS21 | RDX | KCNN2 | LXN | OLFML3 | MT-ND4 |
| POLR2L | CALCRL | SOX4 | BLVRB | TNFSF10 | MT-CO3 |
| NRP2 | HTRA1 | EXOC3L2 | MLLT11 | ADM | CHST1 |
| ITM2B | F2R | CD59 | RPL8 | TAGLN | SH3BP5 |
| HES1 | PDLIM1 | GIMAP8 | CREM | FAM43A | SERPINB4 |
| HTRA1 | LDB2 | HEY2 | HOXB6 | PXDN | MVD |
| NDUFA1 | PBX1 | CDH13 | RPS5 | SNHG25 | ACLY |
| APLNR | TFPI | GSTO1 | SCD | EIF2B2 | SMAD1 |
| COX7C | TNNI1 | THY1 | CBLN1 | RPL22L1 | SMS |
| SEC62 | MYL4 | ESAM | RPL3 | BIRC5 | HSPD1 |
| LBH | TNNT2 | RGS10 | MT-ND1 | LXN | RPL12 |
| RPL34 | LIFR | ITM2B | PRND | NETO2 | FADS3 |
| ENG | TUBB2A | MFGE8 | PCBD1 | ADA | FAM26D |
| MYL7 | NPR3 | FCN3 | NME4 | GALNT1 | ACAT2 |
| C4orf48 | BMP4 | AGRN | MT-ND2 | FASN | RPL18A |
| ATP5J2 | TMEM204 | A2M | GALNT1 | FADS3 | ADAM19 |
| MYEOV2 | GSN | WBP5 | RPS3A | SERPINB3 | GALNT1 |
| PET100 | IVNS1ABP | CAPG | MPC1 | MVD | FAM89A |
| TFPI2 | IL32 | SDCBP | RPS9 | IER3 | C10orf10 |
| FCGRT | EFNB2 | PDLIM1 | C14orf1 | RNF19A | SERPINB3 |

| | | | | | |
|---|---|---|---|---|---|
| COL15A1 | MGST3 | SPP1 | MT-ATP6 | FADS2 | FAM213A |
| OST4 | NRP2 | CD55 | ALDOA | SQLE | CAV1 |
| RPS26 | SOX4 | CLEC14A | TM7SF2 | S100A4 | PHLDA1 |
| TOMM7 | IL6ST | MALAT1 | PSMA2 | WFS1 | MPG |
| VAMP5 | EFEMP1 | TAGLN2 | STRA13 | ICAM2 | GTF3A |
| SPP1 | ENG | TSPAN13 | SORBS2 | HSPA5 | MT-ND1 |
| TMA7 | NPPA | CTNNB1 | EEF1B2 | FADS1 | ICAM2 |
| ATOX1 | TSC22D1 | LIFR | MT-CYB | RP11-322E11.5 | KRT18 |
| RPL36 | COL15A1 | NQO1 | PHLDA2 | RPL26 | ADA |
| UQCR11 | MTUS1 | DUSP4 | MT-ND4 | KLF6 | RPSA |
| C14orf2 | RGS10 | CTSA | RPL7A | LGALS1 | FADS2 |
| ID3 | TSPAN7 | DCHS1 | RPL14 | ITPKB | FAM43A |
| COX6C | IGFBP4 | GIMAP7 | GTF3A | FLNB | RPL22L1 |
| TCEB2 | RBP1 | ZFP36L1 | RPL10 | NFIA-AS2 | SLC12A2 |
| SELE | ADGRL4 | RNF181 | EEF1D | ATF4 | CREM |
| SEC61G | PLPP3 | COL15A1 | RNASE1 | EFCAB14 | TP53I11 |
| CA2 | C11orf96 | EMC10 | RPL10A | RPS2 | MT-ND2 |
| ID2 | EVA1B | HLA-E | CXCR4 | ANXA6 | TUBA1B |
| THY1 | PALMD | ATP1B1 | RPL13 | EDN1 | ACKR3 |
| UQCRQ | HEY2 | RAMP2 | FAM43A | IGF2 | EBP |
| PIN4 | GJA4 | BMP4 | SRM | MGLL | SSR3 |
| CLEC14A | SDCBP | MGST3 | ACAT2 | TMEM97 | MPC1 |
| PLPP3 | HIST1H4C | NFKBIA | RPS4X | RPL36A | |
| CTGF | CD9 | ACTA2 | RPL9 | LY6H | |
| RPL35A | ID2 | FAM107B | EBP | APLN | |
| RDX | S100A16 | JUNB | RPS10 | MT-ATP6 | |
| F2R | C12orf57 | ANKRD1 | GAPDH | TMEM178A | |
| TMSB10 | WBP5 | PLVAP | ADA | FAM26D | |
| NDUFB4 | TUBB2B | CD9 | SERPINB3 | MT-ND6 | |
| TAC1 | MAP1LC3A | IGFBP4 | C20orf24 | ADAMTS7 | |
| RPL41 | ESAM | PALMD | PRELID1 | CAV1 | |
| SNRPG | MMRN1 | DUSP5 | FADS2 | SNHG8 | |
| COMMD6 | CD55 | ID1 | RPL7 | SRM | |
| ITGA1 | DUSP5 | TMEM173 | CHCHD2 | DDX21 | |
| MEF2C | CCDC3 | ELK3 | FAM69B | EBP | |
| KRTCAP2 | COL11A1 | HPGD | PPDPF | SOX18 | |
| SNRPF | LPAR6 | LIPG | YBX1 | RGS20 | |
| SHFM1 | SLC2A3 | MDK | FADS1 | ACAT2 | |
| A2M | MYL12B | MRPL17 | TUBA1B | PCBP4 | |
| THBS1 | MALAT1 | MYL7 | PGP | HMGCS1 | |
| GSTO1 | TFPI2 | ADAMTS9 | RPL13A | RPSA | |
| ADGRL4 | GJA5 | LDB2 | EIF4EBP1 | MT-ND4 | |
| NDUFC1 | EFEMP2 | EFEMP2 | EIF2B2 | CRELD2 | |
| RPS27L | TGFB1 | EFNB2 | CLU | ANGPT2 | |
| COX7A2 | CAPG | CALCRL | CLTB | RPLP0 | |
| MIR4435-2HG | HIPK2 | TFPI2 | OAZ1 | HOMER3 | |

| | | | | |
|---|---|---|---|---|
| CDH13 | NFKBIA | TSPAN7 | PPP1R14B | MTDH |
| COX7B | NGFRAP1 | F2R | RPL4 | TK1 |
| HIST1H4C | OAZ2 | VAMP5 | CAV1 | PCDH12 |
| NAA38 | LMO2 | RNF145 | RPL12 | MYH10 |
| ATP1B1 | CD151 | MT2A | RPL18A | FSCN1 |
| SMIM4 | TPM1 | HIST1H4C | TUBA4A | COX20 |
| TIMM8B | VAMP8 | S100A16 | AURKAIP1 | RPS21 |
| CALCRL | WWTR1 | LBH | RPL19 | CXCR4 |
| RPL35 | OCIAD2 | C12orf57 | NQO2 | TSTA3 |
| UQCR10 | SLC45A4 | TMEM88 | MAP1A | STK32C |
| FLT1 | HSPG2 | TIMP1 | BIN1 | DAAM1 |
| AC090498.1 | RAB31 | ACTC1 | HLX | MTHFD2 |
| MRPL33 | IER2 | NRP2 | MT-CO3 | MPC1 |
| RPL31 | DCHS1 | CA2 | FADS3 | SEPW1 |
| CD9 | LIMD2 | RAB31 | JTB | MT-ND2 |
| EFEMP2 | CLEC14A | IL6ST | RPS19BP1 | BOK |
| COX17 | MEF2C | ITGA1 | ABCA9-AS1 | USP31 |
| ITGB1 | SNRK | FAM101B | MPG | CCND2 |
| COX5B | HLA-E | MDFI | ICAM2 | PTTG1 |
| NOP10 | MT2A | APLNR | ACKR3 | TACC1 |
| LIPG | CST3 | HSBP1 | CREB3L1 | ESYT1 |
| UQCRB | CYYR1 | GRN | SERP1 | DRC1 |
| GSN | PLN | ITGB5 | SH3BP5 | TRIM16 |
| COX6B1 | CTSC | TNNC1 | GPM6B | SDF2L1 |
| KTN1 | FAM107B | HTRA1 | APRT | PRND |
| NDUFB2 | SQSTM1 | TMEM37 | RPL5 | ANXA3 |
| MT2A | IGFBP7 | ID2 | CXorf36 | WSCD1 |
| COTL1 | ATP6V1G1 | APP | VIM | LRRFIP1 |
| SYMPK | ATP1B1 | SELE | SH3BGRL3 | AMD1 |
| NFKBIA | SULT1C4 | PBX1 | RPS6 | MYO1B |
| RPL36A | MRPL17 | BSG | SNRPN | GFOD1 |
| ESAM | SNCA | SRPX | ARL6IP4 | MZT2B |
| LINC00152 | MSMP | HAPLN1 | TPT1 | SSH2 |
| NDUFA2 | GIMAP7 | EFEMP1 | PPA1 | RPL41 |
| HYAL2 | HSBP1 | MEF2C | ADAM19 | RPS10 |
| C12orf57 | BSG | TFPI | GFOD1 | RPL12 |
| S100A6 | TIMP3 | CTSC | HOPX | N4BP3 |
| NDUFB3 | TIMP1 | ESM1 | RPL29 | PTPRF |
| BMP4 | LYSMD2 | | MYL6B | LSS |
| GRN | CD99 | | RP11-322E11.5 | KRT18 |
| SNRPE | PNRC1 | | PRDX5 | HN1 |
| A1BG | CTSD | | SLC12A2 | PPP1R14B |

| | | | | |
|---|---|---|---|---|
| TSPAN7 | DUSP4 | | FAM26D | MT-CO3 |
| ADAMTS9 | MYL3 | | PRR13 | FDFT1 |
| DUSP5 | FAM101B | | ITGB1BP1 | MT-ND1 |
| FSTL1 | SELE | | HOMER3 | CREM |
| ATP5L | CNN3 | | RPL18 | GPRIN3 |
| SERF2 | PGLS | | ZFAND2A | MAP1LC3B |
| MRPL52 | PRKCDBP | | MT-CO2 | EBF3 |
| RPLP2 | AGRN | | DESI1 | INTS3 |
| SON | COTL1 | | PRDX6 | CITED4 |
| DPM3 | UBE2J1 | | NACA | CEACAM21 |
| TFPI | SLC9A3R2 | | ACLY | SOX11 |
| DCHS1 | TAX1BP3 | | ALDH2 | GJA1 |
| LDB2 | MYL12A | | SMAD1 | IQSEC3 |
| SGK1 | EXOC3L2 | | UFD1L | SMS |
| RPS15A | ATP6V0E1 | | HSPD1 | MPG |
| SNRPB2 | THY1 | | RPL22L1 | LAMA4 |
| CTSA | SERPINB6 | | TMEM219 | CACNG8 |
| COL6A2 | BCAM | | SSNA1 | EPOR |
| RPL23 | ATP11C | | SMS | GNG2 |
| C8orf59 | TPPP3 | | MRPS15 | FAM213A |
| TXNDC17 | MARCKSL1 | | BTF3 | COL4A2 |
| PBX1 | RHOC | | | MESDC1 |
| RAB31 | MYH7 | | | MT-ND3 |
| CAPG | NQO1 | | | DOCK9 |
| S100A16 | TMEM173 | | | GTF3A |
| EIF5B | SRPX | | | DBI |
| CTSC | ITGA1 | | | MT-CO2 |
| MT-ND4L | SKP1 | | | FAM126A |
| TOMM5 | CTSB | | | MYZAP |
| HEXB | NDUFB5 | | | ACLY |
| RPL27 | AP1S2 | | | INSIG1 |
| RPL22 | FTH1 | | | SLC7A1 |
| LYVE1 | ZNF428 | | | SLC25A1 |
| COX8A | CSRP2 | | | EPS8L1 |
| CD55 | ANKRD1 | | | LDHA |
| DYNC1I2 | RBPMS | | | BCL2L11 |
| DDX18 | TSPAN13 | | | LIMA1 |
| UQCC2 | KLHL4 | | | ADAM19 |
| ZSCAN16-AS1 | CCDC68 | | | MSMO1 |
| SERPINH1 | CTNNB1 | | | BCYRN1 |
| CDH5 | GRB10 | | | HSPD1 |
| PDIA3 | GNG7 | | | SNHG19 |

| | | | | |
|---|---|---|---|---|
| ACTC1 | TMEM100 | | | MTSS1 |
| LSM7 | ADAMTS9 | | | YIPF1 |
| TAGLN2 | TALDO1 | | | RPL18A |
| EFEMP1 | FDX1 | | | HRAS |
| CTNNB1 | CTSA | | | CDC42EP5 |
| MTRNR2L8 | SAT1 | | | SERTAD4-AS1 |
| EIF3J | C11orf31 | | | SCN3B |
| ADGRF5 | EIF4A2 | | | RPL23A |
| MDFI | SLC7A11 | | | UBE2S |
| SPATS2L | JUNB | | | FMNL3 |
| SRPX | CD40 | | | CLDN5 |
| LINC00116 | DAB2 | | | ILDR2 |
| TMEM204 | CSRP3 | | | BCAT1 |
| PDLIM1 | UCHL1 | | | |
| CMTM3 | FXYD5 | | | |
| TMEM256 | VGLL4 | | | |
| TIMP1 | BACE2 | | | |
| S100A13 | RPS27L | | | |
| FAM107B | ARHGDIB | | | |
| ATPIF1 | C8orf4 | | | |
| RAMP2 | ACTA2 | | | |
| APP | SH3BGRL | | | |
| CD59 | LEPROT | | | |
| MAP1LC3A | ARF5 | | | |
| IL6ST | SCP2 | | | |
| ITGB5 | COL6A2 | | | |
| HEY2 | GPX7 | | | |
| CLEC2B | TGM2 | | | |
| SDCBP | BCAP29 | | | |
| IER2 | BEX4 | | | |
| MMP1 | MFGE8 | | | |
| RPS17 | PRDX2 | | | |
| MINOS1 | RAB11A | | | |
| COX6A1 | TAGLN2 | | | |
| IGFBP4 | SIGIRR | | | |
| MGST3 | ESD | | | |
| EXOC3L2 | ELK3 | | | |
| TDG | CTC-526N19.1 | | | |
| LIFR | ASAH1 | | | |
| KDR | KCNN2 | | | |
| AAMDC | LYL1 | | | |
| TNNC1 | TEAD2 | | | |

| | |
|---|---|
| LINC00493 | TGFBR3 |
| PDGFA | APP |
| CTSB | GIMAP8 |
| COX7A1 | ITGB5 |
| TM4SF18 | RNF181 |
| RNF145 | BST1 |
| CMSS1 | GPX4 |
| RPS25 | BLVRB |
| BSG | B2M |
| RPN2 | TMEM37 |
| MALL | PAPSS1 |
| EIF5A | PGRMC1 |
| LPAR6 | ZFP36L1 |
| S100A11 | PLD3 |
| TMEM37 | MAGED2 |
| LPL | SNTB1 |
| PDIA6 | SIPA1 |
| NDUFA6 | GPX8 |
| SREK1IP1 | RNF7 |
| CD81 | SNX3 |
| ATP1B3 | HDAC1 |
| TMEM173 | ARGLU1 |
| CDC42SE1 | PLIN2 |
| ITGA5 | SLC25A4 |
| FXYD6 | RRAGA |
| NPPA | TSPAN18 |
| ITGA2 | PFN2 |
| ITGA8 | EMC10 |
| WBP5 | SSBP4 |
| VIMP | CD63 |
| ACTA2 | CITED2 |
| DUSP4 | GABARAPL2 |
| NQO1 | IFITM3 |
| TM4SF1 | SETBP1 |
| SNRPD2 | VPS28 |
| WWTR1 | FXYD6 |
| MDK | PCTP |
| NDUFA12 | ISYNA1 |
| IFITM3 | APLNR |
| C19orf70 | ANXA2 |
| MALAT1 | MDFI |
| ELK3 | ISCU |

| | |
|---|---|
| PCDH17 | RABEP1 |
| KCNN2 | TMOD2 |
| TMEM88 | INSR |
| MIF | FABP5 |
| EIF2S2 | PSMD4 |
| JUNB | KLHDC8B |
| RGS10 | REPIN1 |
| PALMD | PLA2G16 |
| LAPTM5 | F2RL2 |
| ZFP36L1 | VEGFC |
| UPF2 | PYURF |
| RPL30 | COMMD7 |
| HACD3 | PEPD |
| NDUFS5 | WDR83OS |
| SRRM1 | GRN |
| TSPAN13 | PPIC |
| RPS15 | EHD3 |
| HLA-E | LIPG |
| RBX1 | LINC00998 |
| BAMBI | GDF15 |
| HSBP1 | PSMB1 |
| EMC10 | UPP1 |
| ATP6AP2 | H3F3A |
| GIMAP4 | ROBO1 |
| AKIP1 | CRIP2 |
| ESM1 | RNF145 |
| MRPS21 | RENBP |
| TNFRSF12A | NRP1 |
| ATP6AP1 | ANXA5 |
| VAMP8 | CDH13 |
| LAPTM4A | GPX1 |
| HSP90B1 | TRAF4 |
| BOLA3 | NME3 |
| CD151 | CIB1 |
| NDUFS6 | RNF130 |
| ENC1 | ARMCX6 |
| AGRN | RHOBTB3 |
| MFGE8 | BRI3 |
| METAP2 | TRIOBP |
| FAU | HECW2 |
| SNHG19 | NUDT14 |
| HLA-DRB1 | RELB |

| | |
|---|---|
| COX14 | LRRC4 |
| MAFB | ANO6 |
| SPRY1 | ZNF771 |
| PIEZO2 | SKIL |
| RNF181 | MAGOH |
| PLK2 | ADGRA2 |
| CLEC11A | NTHL1 |
| GIMAP8 | LIPA |
| GIMAP7 | FSTL1 |
| DAB2 | SOD1 |
| MRPL17 | MDH1 |
| NSRP1 | DHRS3 |
| PSAP | UBB |
| FAM101B | EMCN |
| ATP5J | CLIC1 |
| EFNB2 | ERGIC3 |
| | ARL2 |
| | CD59 |
| | VAMP2 |

Table III: Upregulated and downregulated genes in iPS-CMs in coculture.

| UPREGULATED IN COCULTURE IPS-CM | | | DOWNREGULATED IN COCULTURE IPS-CM | | |
|---|---|---|---|---|---|
| **HEL47** | **HEL24_3** | **BOTH** | **HEL24_3** | **HEL47** | **BOTH** |
| MT-ND3 | GNG11 | GFOD1 | PSMA2 | TCAP | HSPB3 |
| ID3 | HES4 | HES4 | RPL36A | FTL | COL21A1 |
| GFOD1 | A2M | DGKI | COL21A1 | PGAM2 | TCAP |
| MMP15 | PLVAP | ID3 | ESD | FTH1 | FTH1 |
| CAPN2 | ESM1 | MASP1 | EIF3E | MTRNR2L8 | FTL |
| MYOCD | EGFL7 | BAMBI | WSB1 | COL21A1 | ETFB |
| MEF2C | S100A10 | S100A10 | PPP1R1A | LINC01405 | IGFBP5 |
| BMP2 | S100A11 | TMEM141 | H3F3B | AC090498.1 | |
| SDC3 | ID3 | C4orf48 | HSPB3 | LRRC10 | |
| GABRB2 | HSPG2 | | EIF3M | HSPB3 | |
| ATP6V0B | ACTB | | RPL26 | FGF18 | |
| NES | TMEM141 | | CIRBP | ATP5I | |
| CALU | USP18 | | ETFB | NREP | |
| ATP1A1 | C4orf48 | | HAND2-AS1 | TNNI3 | |
| BAMBI | HOPX | | LDHB | NDUFB1 | |
| JAG1 | ATF3 | | CKM | FITM1 | |
| UBE2S | MASP1 | | KCNQ1OT1 | ETFB | |
| SPARC | SH3BGRL3 | | PPDPF | NPPB | |
| MYLK3 | TMSB4X | | TUBA1A | MYL2 | |
| AC090498.1 | RAMP2 | | MDK | SLC8A1 | |
| CCDC85B | ID1 | | IGFBP5 | IGFBP5 | |
| RPS29 | ACTG1 | | NPM1 | CCDC34 | |
| TMEM141 | DSP | | ZFP36L1 | HIST1H1C | |
| MIF | KRT8 | | RPS10 | RP11-532N4.2 | |
| VCAN | THBS1 | | LGALS3 | | |
| S100A10 | TMSB10 | | NDUFA13 | | |
| MASP1 | BAMBI | | TCAP | | |
| TGM2 | APLN | | EGR1 | | |
| MTRNR2L8 | PRSS23 | | FTH1 | | |
| NPPA | KRT18 | | FTL | | |
| C4orf48 | MARCKSL1 | | | | |
| LINC00881 | SLC38A2 | | | | |
| H2AFZ | HES1 | | | | |
| HES4 | DGKI | | | | |
| VASH1 | TM4SF1 | | | | |
| DGKI | GFOD1 | | | | |
| ANKRD1 | VAMP5 | | | | |
| HIST1H4C | CD9 | | | | |
| NPPB | LIMS1 | | | | |
| MT-ND3 | ETV5 | | | | |

| |
|---|
| ID3 |

Table IV: BMP-pathway genes expressed at very low or non-detectable levels in iPS-ECs and iPS-CMs (HEL47)

| **BMP-pathway genes with very low or non-detectable expression in iPS-ECs** | **BMP-pathway genes with very low or non-detectable expression in iPS-CMs** |
|---|---|
| ACVR2A | ACVR2A |
| AVCR1C | BMPR1B |
| BMPR1A | ID1 |
| BMPR1B | SMURF2 |
| ACVR1 | ACVR1 |
| BMP10 | BMP10 |
| BMP5 | BMP4 |
| BMP6 | BMP6 |
| BMP7 | BMP8A |
| BMP8A | BMP8B |
| BMP8B | GDF5 |
| GDF5 | GDF6 |
| GDF6 | GDF7 |
| GDF7 | SMAD1 |
| ID4 | SMAD6 |
| SMAD6 | SMAD7 |
| SMAD7 | SMAD9 |
| SMAD9 | SMURF1 |
| SMURF1 | |

Table V: Upregulated and downregulated genes in Flow exposed iPS-ECs. HEL47.2 and HEL24.3 scRNASeq data are presented, alone, together and with bulk RNASeq.

| **UPREGULATED IN FLOW IPS-EC** | | | | **DOWNREGULATED IN FLOW IPS-EC** | | | |
|---|---|---|---|---|---|---|---|
| **HEL47.2 scRNASeq** | **HEL24_3 scRNASeq** | **BOTH scRNASeq** | **BOTH scRNASeq and bulk RNASeq** | **HEL47.2 scRNASeq** | **HEL24_3 scRNASeq** | **BOTH scRNASeq** | **BOTH scRNASeq and bulk RNASeq** |
| IGFBP5 | SLC9A3R2 | A2M | ADAM15 | ANGPT2 | ANGPT2 | ABCA9-AS1 | ANGPT2 |
| SLC9A3R2 | IGFBP5 | ADAM15 | AK8 | TPM2 | EDN1 | ADM | APLN |
| APLNR | NEAT1 | ADAMTS9 | APOLD1 | EDN1 | ESM1 | ALDOA | APRT |
| G6PC3 | FTH1 | AIF1L | ATOX1 | ESM1 | CXCR4 | ANGPT2 | ARHGAP18 |
| CYP1B1 | EMP3 | AK8 | ATP6V1D | APLN | PGF | ANXA1 | ARHGDIB |
| PLVAP | APLNR | APLNR | BCAT1 | PGF | SERPINB4 | ANXA3 | ARHGEF28 |
| S100A6 | HMOX1 | APOE | BCL6B | SERPINB4 | CTGF | APLN | BST1 |
| PODXL | KLF2 | APOLD1 | BRI3 | CTGF | PRSS23 | APRT | CCND2 |
| TSC22D1 | CYP1B1 | ATOX1 | C11orf96 | CAV1 | C10orf10 | ARHGAP18 | CDK2AP2 |
| NQO1 | NQO1 | ATP1A1 | CALCRL | TAGLN | PIM3 | ARHGDIB | CITED2 |
| TIMP3 | TIMP3 | ATP1B3 | CARHSP1 | UACA | LINC01480 | ARHGEF28 | COPS6 |
| KLF2 | FTL | ATP6V1D | CD276 | MYL9 | CYR61 | BST1 | CRACR2B |
| EMP3 | PODXL | BCAT1 | CD34 | RPS2 | ADM | C10orf10 | CRELD2 |
| NRP2 | SLC2A1 | BCL6B | CD81 | HMGA1 | CHST1 | C1QBP | DCTPP1 |
| HYAL2 | PLVAP | BIN1 | COLGALT1 | GPX1 | LXN | CA2 | DCXR |
| HAPLN1 | S100A4 | BMP1 | CYP1A1 | RPL3 | GPX1 | CALD1 | DDAH1 |
| RGS5 | NRP2 | BRI3 | CYP1B1 | FABP5 | TUBA1B | CAV1 | DYNLT1 |
| PMP22 | S100A6 | C11orf96 | CYYR1 | VIM | PHLDA1 | CBLN1 | EDN1 |
| PRCP | HES4 | C12orf57 | DUSP3 | RPLP0 | ARHGDIB | CCDC85B | FAAP20 |
| CD52 | PMP22 | C19orf33 | DYRK2 | NME4 | TNFSF10 | CCND1 | FABP5 |
| RPS28 | OAF | C8orf4 | ELK3 | CYR61 | SOX18 | CCND2 | FAM43A |
| PLPP3 | S100A10 | CALCRL | ENDOD1 | ID3 | FABP5 | CDK2AP2 | GALNT1 |
| MT-ND3 | CARHSP1 | CARHSP1 | EVA1B | ARHGDIB | SERPINB3 | CHCHD10 | GMFG |
| ATOX1 | APOLD1 | CCDC109B | FABP3 | LXN | TCF4 | CHST1 | HOXB5 |
| FTH1 | IL32 | CD109 | FAM167B | CHST1 | ARHGAP18 | CISD3 | HYI |
| ATP5E | ZFP36L1 | CD276 | FTH1 | KRT18 | UACA | CITED2 | IMP3 |
| APOLD1 | HAPLN1 | CD34 | FTL | TSPAN15 | APLN | CLEC2B | MAST4 |
| RPL38 | HYAL2 | CD59 | FXYD5 | SSR3 | CLEC14A | COPS6 | MPG |
| GJA4 | RHOA | CD81 | FXYD6 | PIM3 | ID3 | CRACR2B | MPST |
| FXYD6 | GSTO1 | CDKN1A | GCLM | PDLIM2 | GIMAP4 | CRELD2 | MRPL23 |
| CALCRL | A2M | COLGALT1 | GIT1 | RPS6 | GMFG | CTGF | MRPS34 |
| TSPAN13 | C11orf96 | COTL1 | GPRC5C | NPM1 | RNF19A | CTHRC1 | NME4 |
| RPS29 | THBD | COX14 | HEG1 | RPSA | SMS | CXCR4 | NPM3 |
| UBL5 | ADAM15 | CYP1A1 | HIPK2 | RPS3 | PRND | CYR61 | OGFRL1 |
| VAMP5 | CYP1A1 | CYP1B1 | HMOX1 | RPL10A | HLX | DCTPP1 | PDGFB |
| TSC22D3 | SLC7A11 | CYYR1 | HS3ST1 | RPL7A | PDGFB | DCXR | PHLDA1 |
| NEAT1 | FABP3 | DAB2 | HSPA12B | ANXA1 | FAM26D | DDAH1 | PIM3 |
| NDUFA3 | HSPA1A | DDAH2 | HYAL2 | EEF1B2 | ACAT2 | DOCK9 | PRDX5 |
| FABP3 | HEG1 | DUSP3 | IGFBP5 | MGST1 | NETO2 | DYNLT1 | RALA |
| CARHSP1 | DNAJB1 | DUSP4 | ISG20 | RPL4 | HEY1 | EBP | RPL3 |
| C4orf48 | EVA1B | DYRK2 | ITGA9 | BST1 | EIF4EBP1 | EDN1 | RPL4 |
| RPS27 | PRCP | EIF4EBP2 | KCNN3 | RPS4X | THBS1 | EIF2B2 | SERPINB3 |
| RPL39 | VAMP5 | ELK3 | KLF2 | MMP1 | EIF2B2 | EIF4EBP1 | SERPINB4 |
| COX6C | RGS5 | EMP3 | KLF4 | SORBS2 | IMP3 | ERP29 | SLITRK4 |
| THBD | PPP1R14A | ENDOD1 | LAPTM5 | PRSS23 | FAM43A | ESM1 | STUB1 |
| RPL37A | SDCBP | EVA1B | LMO2 | GPX3 | GJA1 | FAAP20 | TFPI2 |
| FAM167B | PLPP3 | FABP3 | MEF2A | PCBD1 | TUBA1A | FABP5 | THBS1 |
| ATP5I | S100A13 | FAM167B | MYLK | LGALS1 | TSPAN15 | FAM101B | TM7SF2 |
| HEG1 | FAM167B | FLT1 | NDRG1 | NME1 | TXNRD2 | FAM213A | TMSB4X |
| STC1 | BRI3 | FN1 | NOS3 | HNRNPA1 | OGFRL1 | FAM26D | TPM2 |
| SLIRP | CALCRL | FTH1 | NQO1 | C10orf10 | DCTPP1 | FAM43A | TRAPPC1 |
| HES4 | ISG20 | FTL | NRP2 | TSPO | CBLN1 | FAM89A | TSPAN15 |
| C8orf4 | FXYD5 | FURIN | OAF | RPS3A | RALA | FKBP10 | TSPO |
| MYEOV2 | PALM | FXYD5 | PALM | SRM | NME4 | GALNT1 | TXNRD2 |
| SLC7A11 | SMAGP | FXYD6 | PLAU | FAM26D | RGCC | GCSH | UACA |
| USMG5 | MALAT1 | GCLM | PLEKHA1 | CCND1 | MRPL15 | GIMAP4 | ZDHHC12 |

| | | | | | | |
|---|---|---|---|---|---|---|
| DDAH2 | TSC22D3 | GIT1 | PLIN2 | RPL5 | SAT1 | GMFG |
| ROMO1 | DUSP4 | GJA4 | PLK2 | RPL8 | CAV1 | GPX1 |
| MEF2C | DDAH2 | GPRC5C | PLPP3 | EEF1D | CD24 | GPX8 |
| ADAM15 | TAGLN | GSTO1 | PLVAP | RPL12 | SRM | GTF3A |
| RPS21 | FXYD6 | HAPLN1 | PLXND1 | FAM43A | IGFBP3 | HEY1 |
| TMSB10 | TSC22D1 | HEG1 | PMP22 | RPL10 | TMED9 | HLA-C |
| OST4 | RAMP2 | HES4 | PODXL | HOXB6 | SPRY1 | HLX |
| COX7C | TMEM173 | HIPK2 | PRKCZ | RPS5 | ABCA9-AS1 | HOXB5 |
| C12orf57 | SLC2A3 | HMOX1 | PTPRB | APRT | SOD2 | HOXB6 |
| S100A4 | BIN1 | HS3ST1 | PTPRE | TUBA1B | CA2 | HPGD |
| WBP5 | S100A3 | HSPA12B | PTPRG | RPL14 | TFPI2 | HYI |
| SDCBP | ATP1A1 | HTRA1 | RAB11A | TXNRD2 | SMAD1 | ID3 |
| MALAT1 | TUBB2A | HYAL2 | RAB31 | RPL7 | CCND2 | IMP3 |
| EMCN | BAG3 | IGFBP4 | RAMP2 | RGCC | MAGED2 | KCNJ2 |
| SHFM1 | KLF4 | IGFBP5 | RAPGEF1 | RPS9 | SLITRK4 | LINC00176 |
| TMEM258 | BGN | IL32 | RAPGEF5 | TCF4 | TM7SF2 | LXN |
| POLR2L | RAB11A | ISG20 | RBPMS | C1QBP | MAP2K1 | MAST4 |
| NOP10 | PPDPF | ITGA9 | RHOA | RBM3 | NCL | MGMT |
| MESDC1 | CCND3 | JUND | RIN2 | ARPC3 | CRELD2 | MGST1 |
| SMAGP | RHOB | KCNN3 | ROBO1 | ALDOA | RBM3 | MPG |
| UQCR11 | TSPAN13 | KITLG | RYBP | EIF4EBP1 | BASP1 | MPST |
| RPL41 | TNXB | KLF2 | SH3PXD2A | PFN1 | BST1 | MRPL15 |
| NDUFB4 | LMO2 | KLF4 | SLC2A1 | NETO2 | APLP2 | MRPL23 |
| NDUFA1 | RAB31 | LAPTM5 | SLC2A3 | GNB2L1 | YIPF1 | MRPS34 |
| ATP5J2 | C19orf33 | LMO2 | SLC6A6 | CLU | CITED2 | NETO2 |
| RPL37 | PTMS | LRRC32 | SLC7A11 | RPL18A | MPG | NME1 |
| SEC61G | SVIP | LTBP1 | SLC9A3R2 | TRAPPC1 | DBN1 | NME4 |
| TUBB2A | CD34 | MALAT1 | SMAGP | PHLDA1 | CTHRC1 | NPM1 |
| PET100 | MESDC1 | MAP3K11 | SPTBN1 | ENO1 | SRSF2 | NPM3 |
| OAF | SERPINE2 | MDK | SSH1 | ALDH2 | PDLIM2 | OGFRL1 |
| PTP4A3 | CYYR1 | MEF2A | STARD4 | RPL13A | DYNLT1 | PDGFB |
| RAPGEF5 | NT5E | MEF2C | TEK | SERPINB3 | TM4SF1 | PDLIM2 |
| S100A13 | CD59 | MESDC1 | THBD | FAM89A | EIF4A1 | PFKP |
| SVIP | UBR4 | MLLT4 | TIMP3 | RPL6 | SSR3 | PFN1 |
| MRPL33 | HSPH1 | MSN | TJP2 | RPL19 | HOXB5 | PGF |
| BCAT1 | RAPGEF5 | MTRF1L | TMBIM1 | RPS4Y1 | CLEC2B | PHACTR2 |
| NDUFB1 | APOE | MYLK | TMEM173 | TM7SF2 | SLC43A3 | PHLDA1 |
| CD276 | SPP1 | NDRG1 | TMEM184B | GALNT1 | RAI14 | PIM3 |
| RPL34 | SDPR | NDUFA12 | TNXB | FAAP20 | CALM1 | PLOD1 |
| SNRPG | TEK | NEAT1 | TSC22D1 | EFEMP1 | RP11-322E11.5 | PRDM1 |
| S100A11 | CDKN1A | NOS3 | TSC22D3 | HOPX | HMGB1 | PRDX4 |
| MDK | S100P | NQO1 | TSPAN13 | THBS1 | RANBP1 | PRDX5 |
| RHOA | MDK | NRP2 | TUBA4A | RPL13 | PRDM1 | PRSS23 |
| CD81 | NDRG1 | OAF | TUBB2A | CCND2 | KCNJ2 | RALA |
| TCEB2 | HIPK2 | PALM | UBR4 | LDHA | SLC4A7 | RAN |
| EVA1B | BCAT1 | PCAT19 | | GPX8 | EIF5A | RANBP1 |
| LMO2 | IGF2 | PEA15 | | IMPDH2 | HPGD | RBM3 |
| TIE1 | NDUFA12 | PLAU | | RPL9 | TRAPPC1 | RGCC |
| NDUFA12 | HSP90AA1 | PLEKHA1 | | GCSH | TPM2 | RGS20 |
| RPS26 | CD276 | PLIN2 | | PDGFB | SLC12A2 | RP11-322E11.5 |
| SLC2A1 | STARD4 | PLK2 | | CHCHD10 | ACKR3 | RPL22L1 |
| COX14 | RRAS | PLPP3 | | GMFG | HIF1A | RPL3 |
| RAB31 | PTGR1 | PLVAP | | TNFRSF12A | GALNT1 | RPL4 |
| ESAM | RHOC | PLXND1 | | RPL15 | EHD4 | RPS19BP1 |
| C11orf96 | MKNK2 | PMP22 | | FAM101B | PDE4B | SAT1 |
| PLAU | PLK2 | PODXL | | SLC25A5 | MYDGF | SERPINB3 |
| SLC2A3 | RP11-14N7.2 | PPP1R14A | | COPS6 | PHACTR2 | SERPINB4 |
| C14orf2 | GCLM | PRCP | | EIF3F | APRT | SLC38A1 |
| MEF2A | HSPA12B | PRKCZ | | HOXB5 | CDK17 | SLITRK4 |
| TMA7 | TMBIM1 | PTMS | | GTF3A | FAM89A | SMS |
| COX7B | HS3ST1 | PTPRB | | NACA | POMP | SOD2 |
| MMRN2 | LRRC32 | PTPRE | | CA2 | SOCS3 | SRM |

| | | | | | | |
|---|---|---|---|---|---|---|
| YWHAH | C8orf4 | PTPRG | | RPL22L1 | SLC38A1 | SSR3 |
| UQCRQ | DYRK2 | QKI | | PYCARD | SRSF7 | STUB1 |
| TMEM173 | ROBO1 | RAB11A | | SOD2 | COPS6 | SYNGR2 |
| LBH | DAB2 | RAB31 | | SMS | EBP | TCF4 |
| HMOX1 | DUSP3 | RAMP2 | | RPL29 | TMEM97 | TFPI2 |
| INSR | GIT1 | RAPGEF1 | | SLC25A6 | TK1 | THBS1 |
| GSTO1 | COTL1 | RAPGEF5 | | IMP3 | RRBP1 | TK1 |
| PIN4 | NAA10 | RBPMS | | CHCHD2 | SH3BP5 | TM4SF1 |
| ROBO1 | ADAMTS9 | RGS5 | | SEPW1 | CCND1 | TM4SF18 |
| LTBP1 | MEF2C | RHOA | | EIF3E | NME1 | TM7SF2 |
| IER2 | PKM | RHOB | | TNFSF10 | MBOAT7 | TMSB4X |
| BCL6B | SQSTM1 | RHOC | | RPS7 | B2M | TNFRSF12A |
| COTL1 | GJA4 | RIN2 | | NPM3 | LAPTM4A | TNFSF10 |
| LYVE1 | PLAU | ROBO1 | | GIMAP4 | PLOD1 | TPM2 |
| PALM | PTPRB | RP11-14N7.2 | | HLA-C | RGS20 | TRAPPC1 |
| ELK3 | ORAI1 | RP11-923I11.8 | | ARHGAP18 | CLDN5 | TSPAN15 |
| SNRPF | JUND | RYBP | | ZDHHC12 | MPC1 | TSPO |
| OSBPL8 | SYNPO | S100A10 | | PRDX6 | SYNGR2 | TSTD1 |
| COX7A2 | PEA15 | S100A13 | | BTF3 | TIMM17A | TUBA1B |
| NDUFC1 | YWHAG | S100A3 | | PLOD1 | THRAP3 | TXNRD2 |
| THY1 | SH3BGRL3 | S100A4 | | HYI | SRSF3 | UACA |
| RPL36 | ATP1B3 | S100A6 | | RPL18 | JUN | YIPF1 |
| UQCR10 | BCL6B | SCARB2 | | EIF2B2 | FILIP1 | ZDHHC12 |
| NDUFB2 | SFRP1 | SDCBP | | CD151 | PRMT1 | |
| SEC62 | RYBP | SH3PXD2A | | CBLN1 | ARHGDIA | |
| TOMM7 | MEF2A | SHE | | NAP1L1 | FUS | |
| IFITM3 | HTRA1 | SLC2A1 | | ISG15 | ROBO4 | |
| PMEPA1 | EPAS1 | SLC2A3 | | MPG | FAM213A | |
| RAMP2 | COX14 | SLC3A2 | | MT-CO3 | FAAP20 | |
| TIMM8B | ITPR3 | SLC6A6 | | NT5DC2 | MYH10 | |
| COX6B1 | PLIN2 | SLC7A11 | | TMEM88 | MYL12A | |
| RNASE1 | KITLG | SLC9A3R2 | | CDK2AP2 | TMSB4X | |
| COX5B | NQO2 | SMAGP | | CITED2 | C1QBP | |
| SERF2 | TUBA4A | SPTBN1 | | VDAC2 | MRPL23 | |
| COX17 | LAPTM5 | SSH1 | | NME3 | TXNDC17 | |
| EDNRB | IGFBP4 | STARD4 | | YBX3 | NRP1 | |
| TMEM2 | PTPRG | SVIP | | HSPD1 | PDHB | |
| TEK | MYLK | SYNPO | | GAPDH | EID1 | |
| HSPA12B | TMEM2 | TEK | | SLC25A3 | EBNA1BP2 | |
| PEA15 | ELK3 | THBD | | AHCY | TM4SF18 | |
| EFNB2 | ADCY4 | TIMP3 | | CISD3 | FDPS | |
| PPP1R14A | PLXND1 | TJP2 | | TSTD1 | RPL22L1 | |
| TNXB | TPT1 | TMBIM1 | | EEF1A1 | HOXB6 | |
| HIPK2 | PTPRE | TMEM173 | | CCT3 | GTF3A | |
| KLF4 | GAPDH | TMEM184B | | C12orf75 | ARF4 | |
| TMBIM1 | HSPB8 | TMEM2 | | SDF2L1 | SSR2 | |
| BRI3 | CCDC109B | TNXB | | NIT2 | PPP4C | |
| OAZ2 | ITGA9 | TSC22D1 | | EEF2 | TNFRSF12A | |
| RP11-14N7.2 | ENDOD1 | TSC22D3 | | YIPF1 | PRDX4 | |
| IL32 | RIN2 | TSPAN13 | | PRDX4 | ANXA3 | |
| ATP1A1 | COLGALT1 | TUBA4A | | RAC3 | MSMP | |
| RPL36A | PRDX6 | TUBB2A | | MPST | CIRBP | |
| PLEKHA1 | RAPGEF1 | UBR4 | | RPS19BP1 | PRDX5 | |
| CALU | KCNN3 | VAMP5 | | CCDC85B | MRPL17 | |
| PNP | DNAJC1 | VAMP8 | | EIF3L | COPB1 | |
| NPW | FN1 | ZFP36L1 | | GSTP1 | ILF2 | |
| MARCKSL1 | RSRP1 | | | RAN | ABCE1 | |
| CD34 | C12orf57 | | | CRELD2 | MORF4L2 | |
| COLGALT1 | CD81 | | | ARL6IP4 | SQLE | |
| PLXND1 | BMP1 | | | POU2F2 | NIFK | |
| PTPRB | PCAT19 | | | ATP1B1 | PSIP1 | |
| FLT1 | RP11-923I11.8 | | | DCTPP1 | HNRNPC | |
| RPS27L | GPRC5C | | | STRA13 | HNRNPD | |

| | | | | | |
|---|---|---|---|---|---|
| HSPE1 | MAP3K11 | | | PABPC1 | VPS25 |
| HMGCS1 | ATP6V1D | | | OGFRL1 | CDC42EP3 |
| CDH5 | ATOX1 | | | APOA1BP | PEG10 |
| PLAUR | PLEKHA1 | | | PARK7 | CCDC85B |
| ADAMTS9 | MLLT4 | | | HACD1 | TMEM98 |
| PLK2 | HSPB1 | | | PEBP1 | TSPO |
| NDUFA13 | ZFAND2A | | | MGMT | DAAM1 |
| RPL35A | UBE2F | | | IER3 | MGMT |
| COMMD6 | HERPUD1 | | | LY6E | IDH1 |
| RHOC | SERPINB1 | | | MT-CO2 | DIAPH2 |
| DUSP4 | MTRNR2L8 | | | PHACTR2 | HYI |
| NOS3 | SEPT9 | | | PRELID1 | ITM2A |
| MMP14 | EHD2 | | | MDH2 | GPX8 |
| ETS1 | RBPMS | | | DDAH1 | HLA-C |
| A2M | CD109 | | | DCXR | DLL4 |
| VEGFC | SLC6A6 | | | RALA | CISD3 |
| CYP1A1 | QKI | | | PPIB | HNRNPAB |
| CD46 | SOD1 | | | TMSB4X | TPM3 |
| GRN | PRKCZ | | | GLTSCR2 | DOCK9 |
| SPAG9 | SLC3A2 | | | SLITRK4 | LEPROT |
| RAB11A | UPP1 | | | TPT1 | EIF4A3 |
| SPTBN1 | VAMP8 | | | GPM6B | FAM198B |
| ISG20 | SHE | | | NHP2 | AFMID |
| FN1 | TJP2 | | | RANBP1 | NGFRAP1 |
| ELOVL1 | TMEM184B | | | C6orf48 | PFKP |
| MT-ND4L | PPM1F | | | STUB1 | XRCC5 |
| BAALC | FAM129B | | | HLX | HNRNPDL |
| PSAP | GLRX | | | MT-CYB | JAK1 |
| ITGA1 | FLT1 | | | EBP | HLA-A |
| PRKCZ | IER5 | | | PTPRF | PGK1 |
| KLHDC3 | SH2B3 | | | EBPL | LINC00176 |
| MGST3 | AIF1L | | | RAB34 | DCXR |
| GIT1 | AK8 | | | PRDX2 | RGS3 |
| ATP5L | HTRA3 | | | MTHFD2 | MGST1 |
| NID1 | SEPN1 | | | CLPP | CALD1 |
| RHOB | ASAP1 | | | EIF3H | PUF60 |
| FCGRT | MTRF1L | | | ARHGEF28 | GCSH |
| SOX4 | SSH1 | | | BLVRB | CRACR2B |
| F2R | LTBP1 | | | SAT1 | CHCHD10 |
| SMTN | TESC | | | C20orf27 | ERH |
| S100A10 | SEPT6 | | | LOXL1 | SRI |
| SON | EIF4EBP2 | | | HPGD | TM2D2 |
| DPM3 | ARHGAP23 | | | OLA1 | MPST |
| ENDOD1 | FURIN | | | RGS10 | ZFAND6 |
| CFDP1 | NOS3 | | | C16orf13 | RPL4 |
| TAGLN2 | SCARB2 | | | SNRPN | MVD |
| JUND | SH3PXD2A | | | MRPL23 | PWP1 |
| PTPRG | PDLIM4 | | | MRPS34 | MARCKS |
| RPL35 | MSN | | | ACTB | SNRPB |
| NDUFA6 | SPTBN1 | | | UFD1L | VIMP |
| BCAM | | | | CRACR2B | NPM1 |
| STARD4 | | | | CXCR4 | HNRNPR |
| SLC3A2 | | | | ATP5A1 | TSPAN4 |
| GNG5 | | | | TRIP6 | WFS1 |
| COL15A1 | | | | SLC38A1 | SF3B6 |
| NDUFB3 | | | | EIF3D | EPS8L1 |
| RENBP | | | | PPA1 | RPL3 |
| SNRPE | | | | PRDX5 | KNOP1 |
| A1BG | | | | NT5C3B | OIT3 |
| RBPMS | | | | ANKRD1 | MAST4 |
| LAPTM5 | | | | EIF3G | PFN1 |
| NAA38 | | | | TMED3 | BIRC5 |
| MGAT1 | | | | SNRNP25 | DARS |

| | | | | | |
|---|---|---|---|---|---|
| VAMP8 | | | | BNIP3 | TMED2 |
| KTN1 | | | | PALMD | NPM3 |
| RPLP2 | | | | MRPL15 | ZDHHC12 |
| MINOS1 | | | | MT-CO1 | ANXA1 |
| CRMP1 | | | | KCNJ2 | CLEC1A |
| HTRA1 | | | | MYL12B | BCLAF1 |
| FTL | | | | GPX4 | RSL1D1 |
| PTPN12 | | | | FADS3 | PDXK |
| APOE | | | | SLC2A4RG | YWHAB |
| CDC42EP2 | | | | TUFM | TSTD1 |
| IGFBP4 | | | | CXADR | ETV4 |
| RPL31 | | | | ATP5C1 | HNRNPF |
| KITLG | | | | FLNB | BCAP31 |
| KDR | | | | TM4SF1 | LYPLA1 |
| UTRN | | | | RPS8 | EMG1 |
| MALL | | | | CTHRC1 | ARPC5 |
| FCN3 | | | | PFKP | RPS19BP1 |
| NDUFA2 | | | | MYL6B | POLR2G |
| ETS2 | | | | PROCR | ITM2C |
| LRRC32 | | | | TXN2 | DDX52 |
| SLC9A3R1 | | | | FKBP10 | FKBP1A |
| ATP1B3 | | | | FKBP8 | PAFAH1B3 |
| SCN3B | | | | TAF9 | MRPL3 |
| HIST1H4C | | | | RPL11 | CDK2AP2 |
| AK8 | | | | GSTK1 | FAM101B |
| ITGA9 | | | | GRHPR | MRPS34 |
| SYMPK | | | | COMT | SCAND1 |
| UNC13A | | | | ANXA3 | DTYMK |
| SYNPO | | | | FIBP | FJX1 |
| DUSP3 | | | | DESI1 | HNRNPA2B1 |
| GCLM | | | | KIF5B | YIF1A |
| MLLT4 | | | | EIF3M | PSAT1 |
| ITGA5 | | | | PGAM1 | TPI1 |
| ARRDC3 | | | | SNF8 | HNRNPM |
| TJP2 | | | | ANP32B | HSPA5 |
| C1orf54 | | | | PSMD13 | YIPF3 |
| LIMS1 | | | | CALD1 | SOX17 |
| SREK1IP1 | | | | BABAM1 | ELOVL6 |
| C8orf59 | | | | ADM | STUB1 |
| TTYH3 | | | | REXO2 | POLD2 |
| BMP1 | | | | MT-ND1 | ALDOA |
| PCAT19 | | | | ID1 | ARHGAP29 |
| PTMA | | | | COX4I1 | MANF |
| CYSTM1 | | | | DKK3 | RAN |
| RBX1 | | | | OXA1L | PDIA6 |
| CYYR1 | | | | MT-ND2 | MRPL22 |
| FRMD4B | | | | ABCA9-AS1 | GPRIN3 |
| RBM39 | | | | MT-ND4 | MAGED1 |
| ATPIF1 | | | | CLEC2B | DDAH1 |
| PTMS | | | | MYC | FASN |
| UQCRB | | | | ATP5D | ARHGEF28 |
| PLEC | | | | ALKBH7 | SEPP1 |
| RIN2 | | | | LINC00176 | PSMC4 |
| MN1 | | | | FAM213A | DUSP6 |
| AIF1L | | | | TPGS2 | C22orf29 |
| PBX1 | | | | LRRC4 | DHCR7 |
| RAP2B | | | | STOML2 | ERP29 |
| DDX18 | | | | TK1 | MORF4L1 |
| CTNNB1 | | | | TMEM219 | ADSL |
| S1PR1 | | | | LIMD2 | SPCS1 |
| VGLL4 | | | | POPDC3 | FKBP10 |
| RYBP | | | | MAST4 | COPE |
| MRPL52 | | | | ITGAE | |

| | | | | |
|---|---|---|---|---|
| RPL36AL | | | | TPM1 |
| TEX264 | | | | DOCK9 |
| DYNC1I2 | | | | GFOD1 |
| TUBB2B | | | | MT-ATP6 |
| PLIN2 | | | | C12orf10 |
| STMN1 | | | | RAB13 |
| EPHB4 | | | | TMEM14C |
| CCDC109B | | | | DYNLT1 |
| IL6ST | | | | IGFBP1 |
| CDC42SE1 | | | | HSPA8 |
| CASKIN2 | | | | UQCRC2 |
| ZFP36L1 | | | | OSTC |
| NDUFS6 | | | | CARD19 |
| GPRC5C | | | | RPS18 |
| RAPGEF1 | | | | CORO1B |
| KRTCAP2 | | | | PSMC5 |
| CDKN1A | | | | PRDM1 |
| KLHL5 | | | | AURKAIP1 |
| RNF181 | | | | PSME2 |
| CCM2L | | | | RP11-322E11.5 |
| BCL2L1 | | | | TP53I3 |
| SOX11 | | | | NFU1 |
| SH3PXD2A | | | | SNRPA |
| COX8A | | | | ERP29 |
| SLFN5 | | | | TXNL4A |
| SEMA4C | | | | JOSD2 |
| NDRG1 | | | | ANXA2 |
| PLEKHB2 | | | | HEY1 |
| SQLE | | | | ACTN1 |
| PCDH12 | | | | CTSZ |
| UNC5B | | | | TFPI2 |
| SH2D3C | | | | GAMT |
| ABI3 | | | | ISOC2 |
| KCNN3 | | | | PLXNA2 |
| TRH | | | | RGS20 |
| CD164 | | | | PIN1 |
| LINC00116 | | | | TM4SF18 |
| FAM65A | | | | SYNGR2 |
| LAMA5 | | | | PLGRKT |
| CD109 | | | | TKT |
| MAP1B | | | | SEC11A |
| UBR4 | | | | P4HB |
| LPAR6 | | | | HCFC1R1 |
| LSM7 | | | | SNCG |
| EBF1 | | | | PCSK1N |
| DAB2 | | | | PGP |
| SHE | | | | ERGIC3 |
| MIDN | | | | |
| CMTM3 | | | | |
| LIPA | | | | |
| COL6A2 | | | | |
| LMNA | | | | |
| KANK3 | | | | |
| QKI | | | | |
| EIF5B | | | | |
| IDI1 | | | | |
| SHANK3 | | | | |
| PTPRM | | | | |
| MAPKAPK2 | | | | |
| FSTL1 | | | | |
| PTPRE | | | | |
| SMIM4 | | | | |
| LDLR | | | | |

| |
|---|
| MRPS36 |
| GLUL |
| RAB7A |
| CALM2 |
| LAMP1 |
| VASH1 |
| HDAC5 |
| ARMCX3 |
| MRPS18C |
| EFNB1 |
| EIF4G2 |
| AQP1 |
| RPS15A |
| NBL1 |
| HS3ST1 |
| KRAS |
| TOMM5 |
| ADAM10 |
| TUBA4A |
| MYLK |
| MIR4435-2HG |
| INPP1 |
| EDIL3 |
| SCD |
| ECM1 |
| ARL4D |
| NR3C1 |
| C19orf70 |
| HSBP1 |
| DCHS1 |
| AMOTL1 |
| SSFA2 |
| KIAA1462 |
| ENY2 |
| MSN |
| TNFAIP1 |
| LINC00152 |
| B4GALT1 |
| SLC14A1 |
| ARHGAP27 |
| AC090498.1 |
| WSB2 |
| MTRF1L |
| C19orf33 |
| SSH1 |
| AQP3 |
| TEAD4 |
| SPATS2L |
| TMEM109 |
| TDG |
| NOTCH1 |
| IGF2R |
| COL1A2 |
| DUSP5 |
| GCH1 |
| KLHDC8B |
| RP11-923I11.8 |
| CAMK1 |
| DNAJA1 |
| TBC1D1 |
| ANXA5 |
| SCARB2 |
| ATP5J |

| |
|---|
| ATP6V1D |
| CD59 |
| COL5A1 |
| RAP1B |
| FRMD8 |
| RPP25 |
| CDKN2D |
| IL3RA |
| LLNLR-245B6.1 |
| LMCD1 |
| CDV3 |
| UQCC2 |
| NDUFA4 |
| TMEM184B |
| ARHGEF15 |
| EIF5A |
| TP53INP2 |
| ARHGEF3 |
| FAM219A |
| RP11-783K16.5 |
| GGT5 |
| NREP |
| ´SEPT6 |
| TRIM2 |
| SNRPB2 |
| CLIC1 |
| AGRN |
| RASIP1 |
| PREX1 |
| NPR1 |
| PDLIM5 |
| BIN1 |
| RPL27 |
| PAM |
| ZFP36L2 |
| FAM127A |
| IFI16 |
| TMEM256 |
| EIF4EBP2 |
| DYRK2 |
| SLC6A6 |
| ZNF467 |
| S100A3 |
| DLGAP4 |
| RHOG |
| EIF3J |
| HLA-E |
| SNHG9 |
| ACTN4 |
| KLF13 |
| ITPRIPL2 |
| STAT6 |
| LSM8 |
| TJP1 |
| CPNE8 |
| LINC00493 |
| KIAA1671 |
| SVBP |
| UBN1 |
| FURIN |
| CFLAR |
| TCN2 |
| JUP |

| |
|---|
| PVRL2 |
| UBALD2 |
| NDUFB7 |
| YES1 |
| ATP5EP2 |
| SASH1 |
| CYFIP1 |
| SEZ6L2 |
| MYCT1 |
| LUZP1 |
| MAP3K11 |
| FXYD5 |

Table VI: Genes that were simultaneously upregulated and downregulated in both coculture and flow exposed HEL47.2 and HEL24.3 iPS-ECs

| **UPREGULATED IN BOTH FLOW AND COCULTURE iPS-EC** | **DOWNREGULATED IN BOTH FLOW AND COCULTURE iPS-EC** |
|---|---|
| A2M | ABCA9-AS1 |
| ADAMTS9 | C10orf10 |
| APLNR | CAV1 |
| C12orf57 | CBLN1 |
| CALCRL | CHST1 |
| CD59 | CXCR4 |
| COTL1 | EBP |
| DAB2 | EIF2B2 |
| DUSP4 | FAM213A |
| ELK3 | FAM26D |
| FLT1 | FAM43A |
| FXYD6 | FAM89A |
| GSTO1 | GALNT1 |
| HAPLN1 | GTF3A |
| HTRA1 | LXN |
| IGFBP4 | MPG |
| MALAT1 | NETO2 |
| MDK | PGF |
| MEF2C | PHLDA1 |
| NQO1 | RP11-322E11.5 |
| NRP2 | RPL22L1 |
| PLPP3 | SERPINB3 |
| PLVAP | SERPINB4 |
| PRCP | SMS |
| RAB31 | SOD2 |
| RAMP2 | SRM |
| SDCBP | SSR3 |
| TMEM173 | TUBA1B |
| TSPAN13 | |
| VAMP5 | |
| VAMP8 | |
| ZFP36L1 | |
